# From bird communities to trophic interactions: successive ecological filters decouple potential and realized insectivory in urban environments

**DOI:** 10.64898/2026.02.25.707889

**Authors:** Laura Schillé, Vanessa Poirier, Frédéric Raspail, Philippe Chaumeil, Pierre Bordenave, Pierre-Alexis Herrault, Alain Paquette

## Abstract

1. Urbanization is widely associated with functional homogenization of ecological communities, including declines in insectivorous bird species and concomitant restructuring of arthropod assemblages. While these changes are often assumed to alter ecosystem functioning through trophic interactions, it remains unclear whether bird insectivory in cities can be inferred from community composition alone, or whether it also depends on which species actively forage, on natural prey availability and on the arthropod guilds exploited.
2. We hypothesized that urbanization (i) restructures both insectivorous bird communities and arthropod prey assemblages, thereby (ii) increasing divergence between trait-based potential and realized insectivory, while (iii) prey use is shaped by natural prey availability and active prey selection.
3. We used 25 plots distributed along a controlled urbanization gradient in Montreal, Canada. We combined acoustic monitoring of insectivorous bird communities, arthropod surveys, and camera-monitored cafeteria-like experiments deployed on 73 trees to quantify trait-based potential insectivory, realized insectivory through direct foraging interactions, natural prey availability and prey selection among three arthropod guilds (lepidopteran larvae, spiders and ants).
4. Urbanization reduced bird functional diversity while also decreasing the availability of high-quality prey, particularly lepidopteran larvae and spiders. Realized insectivory was consistently decoupled from community-based expectations, with actively foraging assemblages systematically less insect-specialized than predicted by broader community compositions (potential insectivory). This mismatch was non-random and only weakly related to urban and vegetation gradients. Prey use was largely independent of natural prey availability and instead reflected consistent preferences for larval prey over other guilds, with some indication that the functional composition of bird foraging assemblages further shaped which prey were exploited.
5. Our results show that realized trophic interactions emerge through successive ecological filters rather than directly from community composition: environmental conditions restructure bird communities and prey availability, subsequent foraging filters determine which species actively participate, and prey selection governs which guilds are consumed. Predicting ecosystem functioning in cities therefore requires considering not only biodiversity patterns but also the ecological processes linking community composition to realized trophic interactions.

**Graphical abstract:** 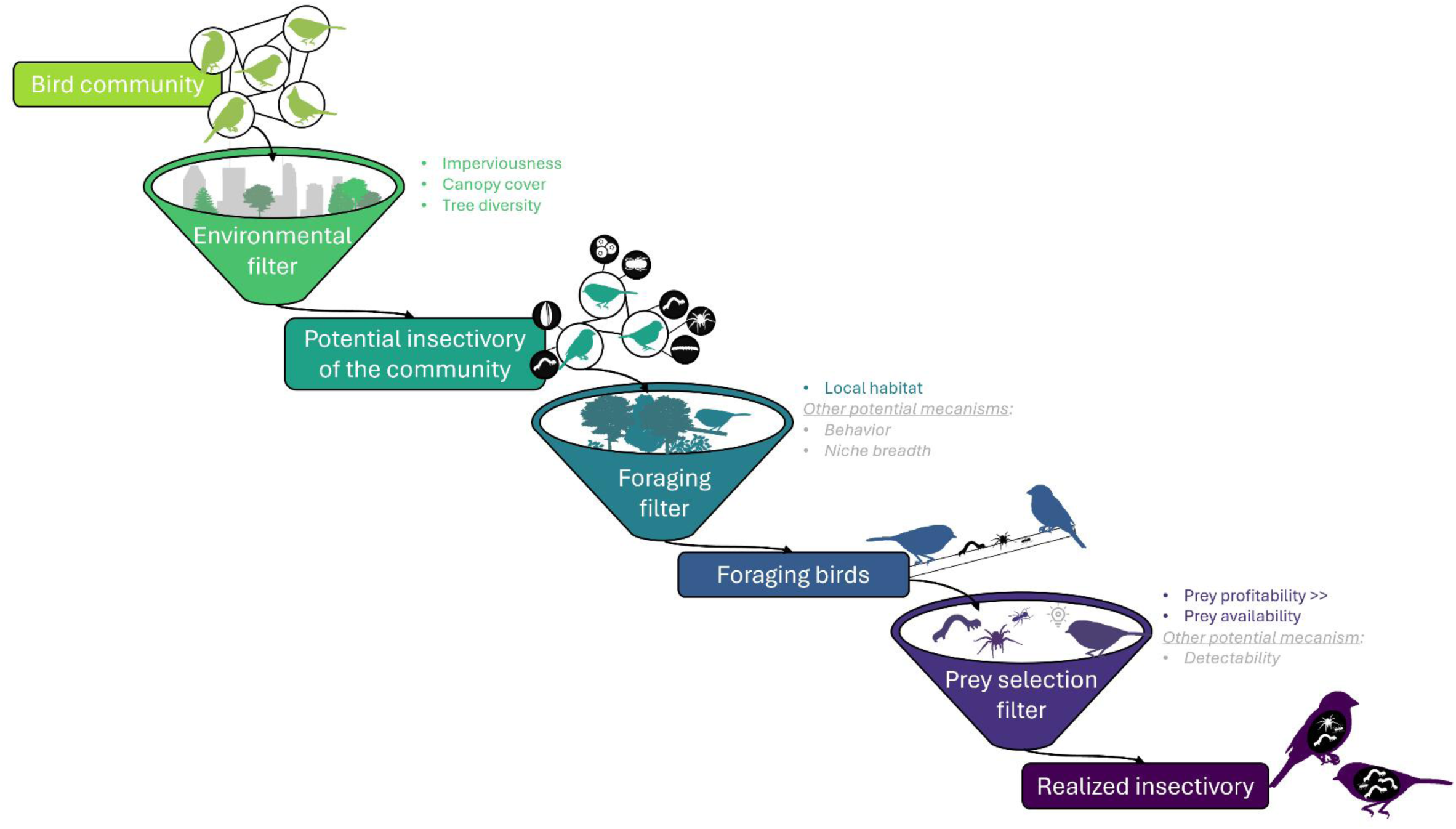

***Conceptual overview of the successive ecological filters linking bird communities to realized insectivory in the city of Montreal, Canada.*** *Environmental filtering first shapes the functional composition of bird communities, defining their potential insectivory. Local foraging conditions then determine which species actively forage, while prey selection governs which prey are ultimately consumed, translating potential into realized insectivory. Mechanisms and predictors shown in grey were not directly tested in this study and are proposed as candidate explanations for the observed patterns.*

## Introduction

Urbanization is reshaping ecosystems at unprecedented rates by altering habitats, resources and disturbance regimes (Alberti et al., 2017; McKinney, 2006). Its effects on biodiversity and community composition, particularly in birds, have been extensively documented (Marzluff, 2001; Sol et al., 2020). However, evidence that these compositional changes alter trophic interactions remains inconsistent (Long & Frank, 2020; Schillé et al., 2025). This gap is critical, as trophic interactions are one of the mechanisms linking biodiversity patterns to ecosystem functioning and services. Urbanization typically filters the biodiversity of bird communities toward a dominance of generalist and disturbance-tolerant species at the expense of specialists (Devictor et al., 2007; Julliard et al., 2006). Such biotic homogenization is frequently assumed to weaken ecosystem functions, including the regulation of arthropod populations (Philpott et al., 2009; Sol et al., 2020). Yet, this assumption has rarely been tested using direct observations of realized bird-arthropod trophic interactions (Palacio et al., 2018).

Instead, predation function provided by birds is typically inferred indirectly from species presence (Melles et al., 2003), relative abundance (Cristaldi et al., 2017), functional traits (Sol et al., 2020), or simplified predation proxies (Long & Frank, 2020). While these approaches provide valuable insights into bird community structure and potential functioning (Palacio et al., 2018; Sol et al., 2020), they implicitly assume that the presence or dominance of insectivorous species reflects their actual contribution to trophic interactions. However, trophic interactions may not solely be determined by community composition. They can vary substantially with local environmental conditions, species interactions, and behavioral flexibility at individual and community levels (Birkhofer et al., 2026; Chatelain et al., 2026; Seress & Liker, 2015; Sinkovics et al., 2021). Thus, bird communities that appear functionally similar based on composition or traits may differ markedly in how they exploit available resources, leading to potential mismatches between potential and realized trophic interactions.

Urban environments provide an ideal setting for investigating these mismatches because habitat structure and resource availability vary considerably across urban landscapes, generating strong environmental filters that may decouple community composition from realized trophic interactions. This environmental heterogeneity encompasses gradients in vegetation structure, anthropogenic disturbance, and prey availability operating across multiple spatial scales (Beninde et al., 2015; Planillo et al., 2021). Urbanization often reduces high-quality arthropod prey such as lepidopteran larvae (Fenoglio et al., 2020), while favoring less preferred but more resilient taxa such as Hymenoptera and Araneae (Magura et al., 2010; Menke et al., 2011). These shifts can alter bird community composition, actively foraging assemblages, foraging behavior, and prey selection, ultimately reshaping realized trophic interactions (Planillo et al., 2021; Sinkovics et al., 2021). Importantly, prey use may not only reflect prey availability, but can also result from active selection processes, particularly under resource limitation or altered prey quality (Nell et al., 2023). Recent work thus emphasizes that understanding the effects of environmental change on food webs and ecosystem functions requires explicitly accounting for resource use and prey selection (Barnagaud et al., 2019; Birkhofer et al., 2026). Overlooking these processes risks biased predictions of the capacity of urban ecosystems to sustain trophic regulation, with direct implications for urban biodiversity conservation and management.

To study how urbanization shapes realized trophic interactions, we directly quantified bird–arthropod prey interactions across a controlled urban gradient using a multi-prey cafeteria-like experiment deployed on 73 trees across 25 plots of the *Montreal Urban Observatory*, Canada (Paquette et al., 2026). Custom-built near-continuous cameras enabled continuous image acquisition, allowing us to identify bird species actually involved in foraging interactions. This approach enabled a direct comparison between trait-based potential insectivory, inferred from the functional composition of the acoustically monitored bird community, and realized insectivory derived from the subset of species detected while actively foraging. We tested whether urbanization (H1) restructures insectivorous bird communities together with arthropod prey availability, (H2) decouples trait-based potential from realized insectivory through differences between community composition and foraging assemblages, and (H3) shapes prey use through the combined effects of natural prey guild availability and active prey selection.

## Material and Methods

### Study design

We conducted the study in June 2025 (the peak breeding season for most insectivorous birds) on the Island of Montreal, Canada (∼2.2 million inhabitants). We used 25 study plots from the *Montreal Urban Observatory* network, distributed along orthogonal gradients of vegetation cover and human population density (**Fig. 1a**). The network included 24 standard plots and one additional highly vegetated plot located in Parc Maisonneuve (plot MNV). Within each plot, we selected three focal trees within a 200 m radius of the plot center (only two trees were available in two of the plots), totaling 73 trees. Selected trees were mature, alive, accessible with a ladder, and located on public land or private land with permission.

**Fig. 1.**
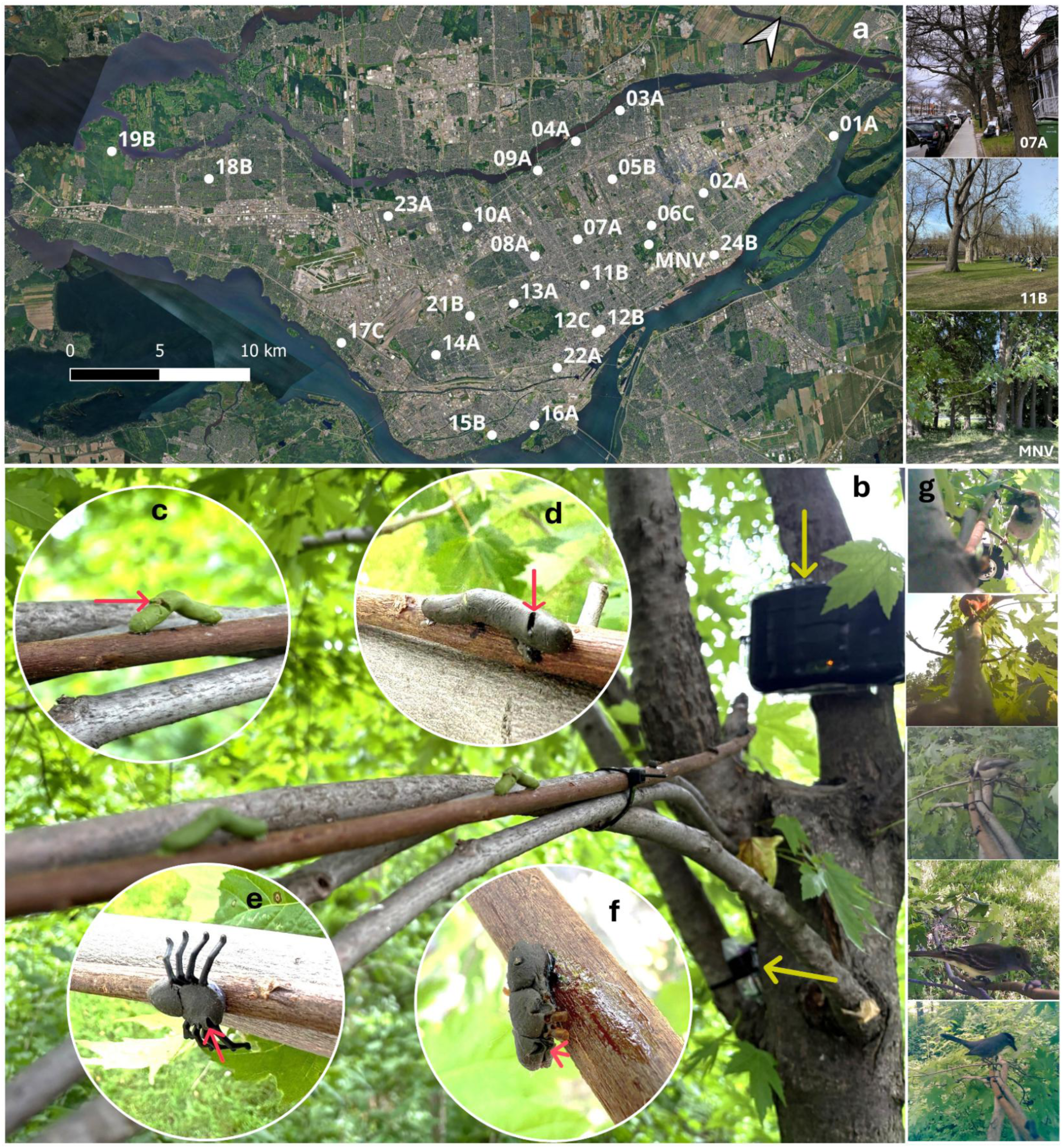
Study design and observed bird attacks on artificial prey. (a) Locations of the 25 sampled urban plots in the city of Montreal with examples of three plots ranging from the most urban (07A, top), to the least urban (MNV, bottom). (b) Audiomoth recorder and custom-built near-continuous camera (yellow arrows) positioned opposite to the willow branch bearing the artificial prey. (c-f) Bird attack marks on artificial prey: (c) green larva, (d) brown larva, (e) spider, (f) ant. (g) Five bird species attacking fake prey, ordered from most urban/tolerant to least (top to bottom): House Sparrow (Passer domesticus), Northern Cardinal (Cardinalis cardinalis), Black-capped Chickadee (Poecile atricapillus), Great Crested Flycatcher (Myiarchus crinitus), Gray Catbird (Dumetella carolinensis).

We focused on silver maple (*Acer saccharinum*), a native, highly abundant species in Montreal known to host diverse arthropod and bird communities (Frank et al., 2013; Gabriel, 1990). When suitable silver maples were not available within a plot, we used Freeman maple (*Acer × freemanii*, a hybrid of *A. saccharinum* and *A. rubrum*) as a substitute (n = 7 trees), randomly distributed across the urban gradient.

### Arthropod prey availability

We sampled arthropod communities on focal trees during the first two weeks of June 2025 (approximately four weeks after budburst). On each tree, we sampled four accessible branches (∼2.5 m height) distributed around the canopy using the beat-sheet sampling method. Each branch was beaten for approximately 10 seconds, and dislodged arthropods were collected with aspirators and preserved in ethanol. Sampling was complemented by 10-minute visual inspections of branches and trunks to collect less mobile arthropods. For all analyses, collected arthropods were pooled at the tree level.

In the laboratory, a single observer identified, counted, and assigned arthropods to four broad prey guilds: lepidopteran larvae (high-quality prey; Sinkovics et al., 2021), spiders (*Araneae*) (nutrient-rich; Ramsay & Houston, 2003), ants (*Hymenoptera*) (less-preferred; Judson & Bennett, 1992; Singer et al., 2017), and others. When ant colonies exceeded 60 individuals, abundance was standardized to 100. For analyses, we retained the total abundance per tree of lepidopteran larvae, spiders, and ants.

### Insectivory of urban bird communities

To bridge the gap between community composition and realized trophic interactions, we quantified insectivory using two complementary approaches: (i) passive acoustic monitoring at the plot scale to characterize the community of functionally insectivorous species (potential insectivory) and (ii) camera-monitored cafeteria-like experiments at the tree scale to identify species assemblages actively foraging on different offered prey (realized insectivory).

### Acoustic identification of bird communities

Bird communities were characterized using autonomous recorders (Audiomoth; Hill et al., 2018), deployed near the center of each plot on 18 June 2025 (six weeks after budburst, corresponding to peak vocal activity and nestling provisioning). Most recorders operated for one week; delays at a few plots (up to three weeks) were deemed negligible given the limited mobility of breeding pairs.

Recordings were processed using automated species identification (BirdNET, Kahl et al., 2021), followed by expert validation by V.P. and filtering to ensure reliable detections (**Appendix S1**). We retained 37 functionally insectivorous (FI) species, defined as species consuming arthropods and foraging in trees during at least part of the year.

At the plot level, we calculated: (i) FI species richness, (ii) FI Shannon diversity, FI functional dispersion (FDis) both weighted by vocal dominance, and (iii) Community-weighted mean (CWM) of the proportion of invertebrates in the diets of FI species, also weighted by vocal dominance (**Eq. S1**). Because arthropods constitute the dominant component of invertebrate prey consumed by FI birds, this dietary metric represents the potential contribution of the local bird community to arthropod consumption based on species composition, hereafter referred to as *potential insectivory*. Trait data (morphological, reproductive and behavioral) were extracted from the Cornell Laboratory of Ornithology (2022) and EltonTraits 1.0 (Wilman et al., 2014) databases (**Table S1**).

### Quantifying realized insectivory using cafeteria-like experiments

On each of the 73 focal trees, we installed a cafeteria-like device on a horizontal branch (∼ 2.5 m above ground) to assess bird prey selection among different artificial prey types. The device consisted of a 150 cm willow twig bearing 12 artificial prey items (**Fig. 1b**): three green caterpillars, three brown caterpillars, three spiders, and three ants (**Fig. 2c-f**). These prey types represent common arthropod guilds consumed by insectivorous birds and differ in nutritional value and urban occurrence. Prey were randomly ordered, spaced 12 cm apart and made of plasticine or plastic models coated with plasticine. Their colors matched those of the corresponding prey observed during arthropod sampling.

**Fig. 2.**
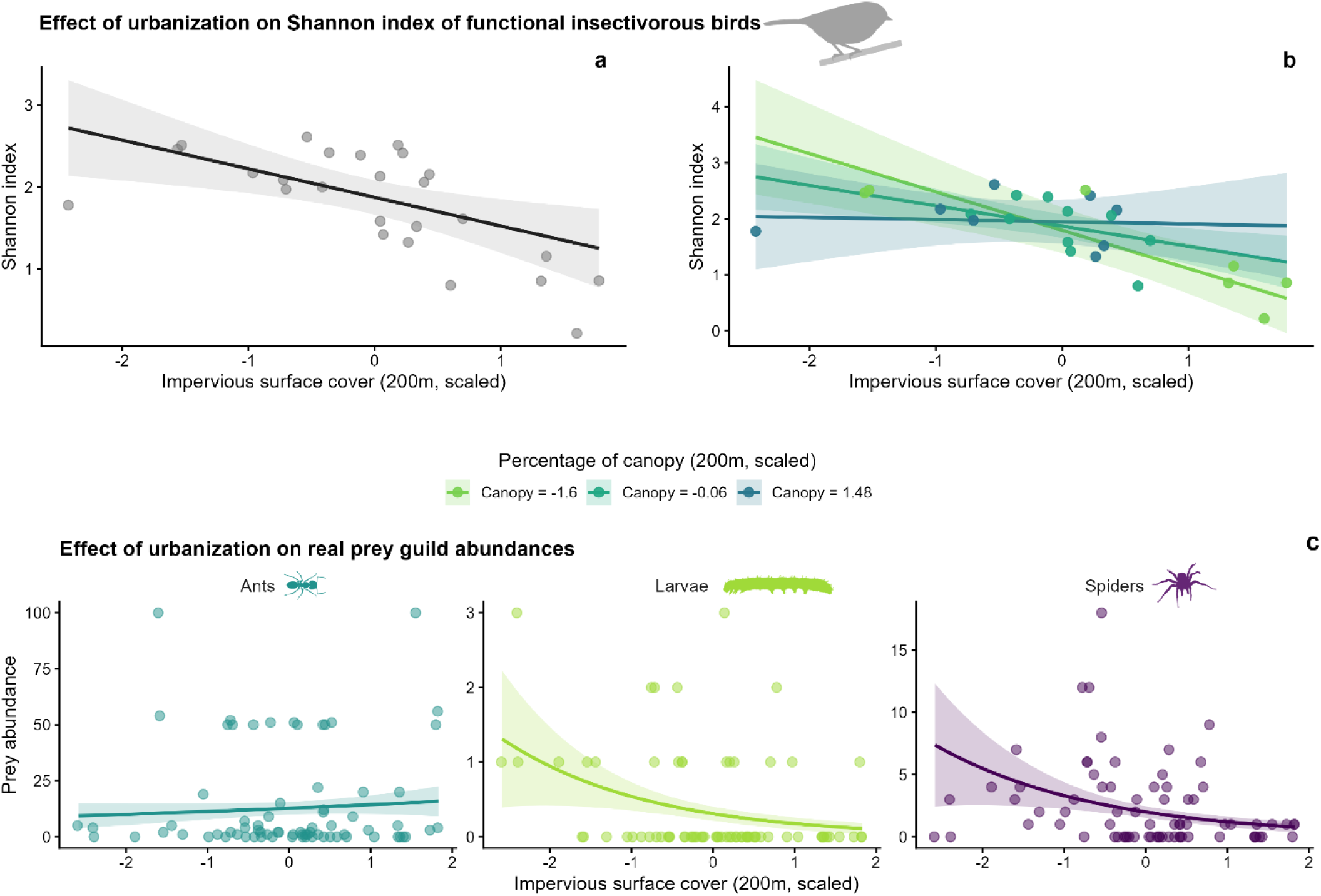
Effects of urbanization on insectivorous birds and the availability of their natural prey. Predicted effects of impervious surface cover alone (a) and in interaction with canopy cover within 200 m (b) on the Shannon diversity index of functional insectivorous birds. Colored lines represent model predictions at low (light green, canopy = –1.6), intermediate (dark green, canopy = –0.06), and high (blue, canopy = 1.48) canopy cover, with shaded areas indicating 95% confidence intervals. Dots show raw data. Observed values for impervious surface ranged from 8.5 to 88.7% within a 200 m buffer; canopy cover ranged from 8.8 to 32.0%. (c) Predicted effects of the interaction between impervious surface cover and prey guilds (ants, larvae, spiders) on natural prey abundance. Shaded areas indicate 95% confidence intervals. Dots show raw data. Observed values for impervious surface ranged from 6.1 to 89.0% within a 200 m buffer.

Cafeteria-like devices were monitored every two days over a 14-day period. At each visit, prey items were scored as attacked (based on bill marks, see **Fig. 1c, d, e, f** for examples) or not, then reshaped, resulting in seven temporal replicates per tree.

To monitor bird interactions with the cafeteria and validate attack marks, we installed a custom-built near-continuous camera (Schillé et al., 2026; **Fig. S1**) on the trunk facing each cafeteria, capturing images at 2 frames per second during daylight hours. Image sequences were stored locally and retrieved every two days.

Images were processed using EcoAssist software with the MegaDetector animal detection model (Beery et al., 2019), using a conservative 2% confidence threshold to minimize false negatives. Detections were clustered and visually verified to retain birds only (**Fig. 2g**).

Bird-cafeteria interactions were classified into four categories: (1) clear attack on an identifiable prey item, (2) active foraging behavior (suspected attack but not clearly identifiable due to limited image resolution or prey being outside the focal area), (3) bird perched on the cafeteria, and (4) bird present, but no visible interaction with the cafeteria prey. For analyses, we retained only categories 1 and 2. Birds were identified to species from images by L.S. We quantified the number of foraging events per species and tree and calculated a CWM of the proportion of invertebrates in species’ diets within the foraging assemblage at the tree level (**Eq. S2**), providing a proxy for *realized insectivory*.

### Environmental predictors

All spatial analyses were conducted in QGIS (v. 3.34). To capture the multidimensional nature of urbanization, we initially selected 12 environmental variables describing habitat structure, anthropogenic pressure, and vegetation characteristics around focal trees (all predictor, aggregation level, definition, resolution, and sources are summarized in **Table S2**). Most predictors were calculated within a 200 m buffer (consistent with typical home ranges of common insectivorous bird species and widely used in urban avian studies; Cornell Laboratory of Ornithology, 2026). Tree diversity metrics were calculated at the plot scale, matching the urban forest inventory resolution. For analyses of realized insectivory and prey selection, we added local vegetation cover within a 20 m buffer around each focal tree to better capture fine-scale habitat features influencing bird foraging behavior (Melles et al., 2003).

To reduce collinearity, the 12 predictors were summarized using a principal component analysis (PCA; **Fig. S2**). The first two axes explained 67.7% of total variance, reflecting gradients of urbanization and tree diversity. We retained four weakly correlated predictors for subsequent analyses: impervious surface (200 m), canopy cover (200 m), plot-level tree Shannon diversity, and local vegetation cover (20 m).

### Statistical analyses

All analyses were conducted in R (v. 4.4.2, R Core Team, 2026). Continuous predictors were scaled before analyses, and model assumptions were checked using standard residual diagnostics. Model significance was assessed using likelihood-ratio tests (LRT), comparing full and null models and, for interactions, nested models with and without the interaction. Where relevant, group-specific slopes and their pairwise comparisons were estimated using the emtrends function (package emmeans), with Tukey-adjusted pairwise comparisons. Full model descriptions are provided in **Table S3**.

### Effects of urbanization on bird communities and prey availability

We first assessed how urbanization shapes (i) the functional structure of insectivorous bird communities at the plot level and (ii) the availability of key arthropod prey guilds at the tree level.

At the plot level, we modeled species richness (Poisson generalized linear model, GLM), Shannon diversity, FDis, and CWM of the proportion of invertebrates in species’ diets within the acoustically-detected insectivorous communities (linear models, LMs) as a function of impervious surface (200m), canopy cover (200m), and plot-level tree diversity.

At the tree level, we modeled the abundance of major prey guilds (lepidopteran larvae, spiders, ants) using zero-inflated negative binomial (ZINB) mixed models, accounting for excess zeros and overdispersion in count data. Fixed effects included prey guild in interaction with an environmental predictor (either impervious surface (200m), canopy cover (200m), local vegetation cover (20m), or plot-level tree diversity), with plot identity as a random intercept. Pairwise post-hoc comparisons among prey guilds were performed using Tukey-adjusted tests.

### Functional mismatch between potential and realized insectivory

To test whether urbanization decouples the functional composition of foraging assemblage from that of the broader present FI community, we compared the CWM of the proportion of invertebrates in the diets of acoustically detected FI species at each plot (potential insectivory) with that of species directly observed foraging at focal trees (realized insectivory).

First, we calculated the functional mismatch between realized and potential insectivory as the difference between the two CWM values. We then tested whether the median mismatch differed from zero using a one-sample Wilcoxon signed-rank test. Second, we assessed whether any observed mismatch could arise from random sampling of the plot species pool. For each tree, null foraging assemblages were generated by randomly assembling species from the plot-level acoustically detected FI community while preserving observed species richness. Species were sampled proportionally to their relative acoustic dominance, thereby accounting for differences in plot occurrence probabilities. Standardized effect sizes (SES) were calculated as: 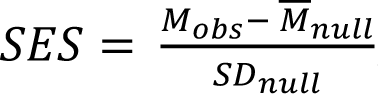, where *M*_*obs*_ is the observed functional mismatch, *M*^⎯^_*null*_ is the mean mismatch expected under the null model, and *SD_null_* is the standard deviation of the null distribution. Negative SES values indicate that foraging assemblages were less insect-specialized than expected under random assembly. Mean SES values were compared against zero using one-sample t-tests.

Finally, to evaluate whether functional mismatch varied along the urbanization gradient, we modeled this mismatch as function of four environmental predictors separately: impervious surface cover (200 m), vegetation cover (20 m), canopy cover (200m), and plot-level tree diversity using generalized linear mixed models (LMMs), with plot identity as a random intercept.

### Drivers of prey use

To understand the mechanisms underlying prey use along the urban gradient, we followed a hierarchical framework. (i) We quantified whether birds exhibited consistent prey preferences among artificial prey types; (ii) we then tested whether these preferences varied across urban environments; (iii) we finally evaluated two non-exclusive mechanisms that could explain variation in prey use: passive tracking of natural prey guild availability, whereby attack rates on artificial prey reflect the relative abundance of corresponding natural prey guilds, and active selection, whereby birds disproportionately attack some prey types relative to their natural availability.

We first tested whether attack probability differed among prey types using a binomial generalized linear mixed model (GLMM) including prey type as a fixed effect. Given the hierarchical experimental design, tree identity nested within plot identity and sampling date were included as random effects.

We then assessed whether attack probability varied along environmental gradients associated with urbanization. Separate binomial GLMMs were fitted including the interaction between prey type and four environmental predictor (impervious surface cover (200m), local vegetation cover (20m), canopy cover (200m) or plot-level tree diversity).

To test whether prey use passively tracked natural prey resource availability, we fitted a binomial GLMM including prey type, relative availability of the corresponding natural prey guild (larvae, spiders or ants), and their interaction. A significant effect of prey availability would indicate that attack probability increases when a prey type becomes locally more abundant.

Finally, we tested whether prey use varied with the functional composition of the foraging bird community. For each plot and prey type, we fitted a binomial GLMM modeling the proportion of attacked prey as a binomial response (attacked vs. unattacked prey) as a function of prey type in interaction with the CWM of the proportion of invertebrates in species’ diets within the camera-observed foraging assemblage. Plot identity was included as a random effect.

Additional analyses examining species-specific foraging patterns were conducted for a subset of common species and are presented as exploratory results in **Fig. S6**.

## Results

To assess how urbanization alters trophic interactions mediated by insectivorous bird communities, we first examined its effects on (i) FI community composition and functional structure and on (ii) prey availability. We then tested for functional mismatches between potential and realized insectivory, and we finally evaluated prey use patterns.

### Urbanization alters the composition of insectivorous bird communities and the availability arthropod prey (H1)

Urbanization strongly affected the composition and functional structure of insectivorous bird communities. Species richness (β ± SE = −0.33 ± 0.10, p < 0.001), Shannon diversity (β = −0.35 ± 0.11, p < 0.01), and FDis (β = −0.037 ± 0.016, p < 0.05) all declined with increasing impervious surface cover (**Fig. 2a** for Shannon diversity; **Fig. S3a** and **Fig. S4a** for species richness and FDis, respectively). These declines were partially mitigated by vegetation structure and composition: the negative effect of impervious surface on species richness was marginally buffered by plot-level tree diversity (**Fig. S3b**; β = 0.21 ± 0.12, p = 0.079; Pseudo-R² = 0.69), while declines in Shannon diversity and FDis were significantly attenuated in areas with higher canopy cover within 200 m (**Fig. 2b**; β = 0.21 ± 0.09, p < 0.05, Adj. R² = 0.46 and **Fig. S4b**; β = 0.034 ± 0.013, p < 0.05, Adj. R² = 0.47, respectively). The plot-level CWM of the proportion of invertebrates in the diets of FI species also decreased along the urban gradient (**Fig. S5**; β = −0.84 ± 0.16, p < 0.001, Adj. R² = 0.57), indicating a convergence toward more generalist diet strategies in highly urbanized environments.

Jointly, natural arthropod prey availability varied significantly among guilds. Lepidopteran larvae were the least abundant prey group, followed by spiders and ants (all pairwise post-hoc comparisons, p < 0.0001). Prey guild abundance also interacted significantly with impervious surface cover (LRT: χ² = 6.45, df = 2, p = 0.040). Along the urbanization gradient, lepidopteran larvae abundance declined significantly with increasing impervious surface cover within 200 m (slope: β = −0.56 ± 0.28, p < 0.044), as did spider abundance (slope: β = −0.51 ± 0.25, p = 0.043), whereas ant abundance did not vary significantly (**Fig. 2c**; marginal R² = 0.49). Models including local vegetation cover, canopy cover (200m), or plot-level tree diversity revealed no significant effects, indicating that impervious surface cover was the dominant driver of variation in natural prey availability.

### Urbanization decouples potential and realized insectivory through non-random filtering (H2)

Across the urban gradient, the functional composition of foraging assemblages consistently differed from that of the broader acoustically detected insectivorous community. The CWM of the proportion of invertebrates in the species’ diets was significantly lower within assemblages actively observed foraging than within the corresponding acoustically detected plot species assemblages (median mismatch = −6.1; Wilcoxon test: V = 377, p < 0.01; **Fig. 3a**), indicating realized trophic interactions were generated by assemblages that were functionally less insect-specialized than expected from community composition alone.

**Fig. 3.**
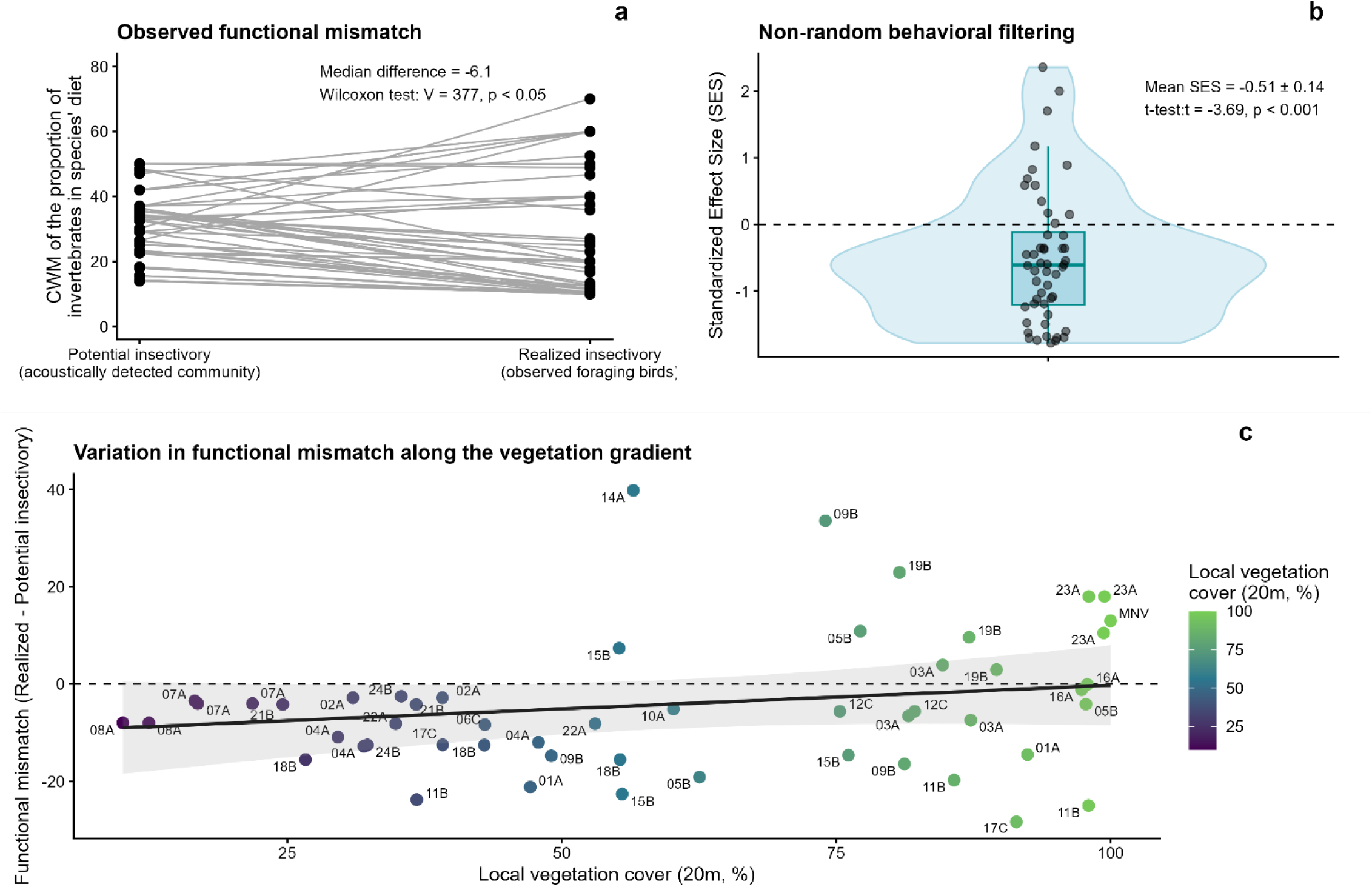
Functional mismatch between potential and realized insectivory. (a) Paired plot showing the CWM of the proportion of invertebrates in species’ diets within the acoustically detected insectivorous community (potential insectivory) and in the observed foraging assemblage (realized insectivory) for each tree. Lines connect paired values. Most lines descend below zero, indicating lower realized insectivory. (b) Distribution of standardized effect sizes (SES) from null model simulations across plots, with the violin showing the density and distribution and the boxplot indicating the the median and interquartile range. Negative SES values indicate that realized communities are less insect-specialized than expected under random assembly. (c) Functional mismatch along local vegetation cover gradient (20 m). The dashed line represents a 1:1 relationship (equality between potential and realized insectivory). Points below this line indicate lower-than-expected realized insectivory given the acoustically detected species pool, while points above indicate higher-than-expected values. Although functional mismatch is not significantly related to vegetation cover, a trend suggests that low-vegetation plots tend to fall below the 1:1 line, whereas highly vegetated plots are closer to or above it, indicating greater agreement between community composition and realized foraging in more locally vegetated habitats.

Null model simulations confirmed that this mismatch did not result from random sampling of the plot species pool. Standardized effect sizes (SES) were significantly lower than zero (mean SES = −0.51 ± 0.14; one-sample t-test: t = −3.70, p < 0.001; **Fig. 3b**), indicating non-random filtering toward less insect-specialized foraging assemblages.

Although the magnitude of functional mismatch was not significantly related to vegetation cover, impervious surface, canopy cover, or tree diversity (all p > 0.05), the mismatch tended to become less negative with increasing local vegetation cover (**Fig. 3c**), from a model-predicted −5.11 (95% CI: −10.39 to 0.17) at 50% vegetation cover to −1.22 (95% CI: −8.22 to 5.78) at 90%. Along this gradient, trees locally surrounded by higher vegetation cover were more frequently associated with levels of realized insectivory consistent with those expected from the acoustically detected plot bird community. In contrast, in sparsely vegetated habitats, foraging assemblages tended to be more functionally decoupled from the plot species pool, being less insect-specialized than expected from community composition.

### Urbanization drives prey use through prey preferences rather than prey availability (H3)

Birds showed marked and consistent differences in prey use across all plots. Attack probability differed significantly among prey types (Likelihood ratio test: χ² = 123.25, df = 3, p < 0.001), with green larvae attacked most frequently, followed by brown larvae, while spiders and ants were attacked least (Tukey pairwise comparisons: all p < 0.01 except spider vs. ant, p=0.79; **Fig. 4a**). This pattern indicates strong prey preferences across the study site.

**Fig. 4.**
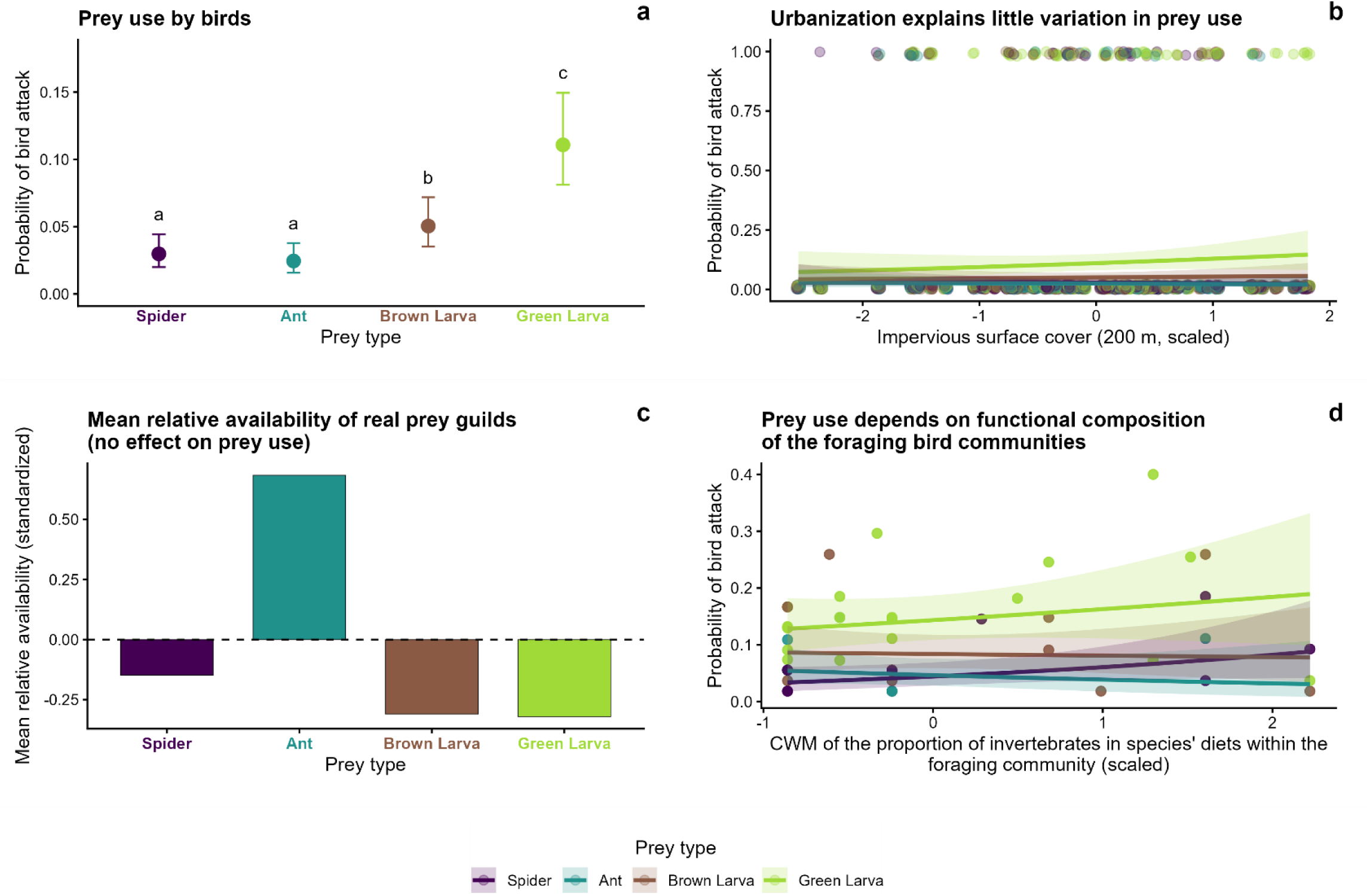
Drivers of prey use by urban bird communities. (a) Predicted attack probability for the four artificial prey types across all study plots. Error bars represent95% confidence intervals and letters indicate significant differences among prey types based on Tukey post-hoc comparisons. (b) Relationship between predicted attack probability and impervious surface cover (200m). The shaded areas indicate 95% confidence intervals. Dots show raw data. Observed values for impervious surface ranged from c.1 to 89.0% within a 200m buffer. (c) Mean relative availability of the natural prey guilds. (d) Relationship between predicted attack probability and the CWM of the proportion of invertebrates in species’ diets within the foraging bird assemblages. The shaded areas indicate95% confidence intervals. Dots show raw data. Observed values for CWM ranged from 10 to c0 within a 20m buffer.

This ranking of preferences remained stable along the urbanization gradient. The interaction between prey type and impervious surface cover was not significant (χ² = 3.85, df = 3, p = 0.28), nor was the interaction with plot-level tree diversity (χ² = 4.33, df = 3, p = 0.23), canopy cover (χ² = 1.66, df = 3, p = 0.65), or local vegetation cover (χ² = 2.51, df = 3, p = 0.47). Accordingly, none of the prey-specific slopes differed significantly from zero for any of these variables (impervious surface: all p ≥ 0.27; tree diversity: all p ≥ 0.13; canopy cover: all p ≥ 0.10; vegetation cover: all p ≥ 0.30), and pairwise comparisons among slopes were also non-significant after Tukey adjustment in every case (all p ≥ 0.18; **Fig. 4b**). Overall, these environmental predictors explained only a small proportion of variation in prey use (marginal R² = 0.09-0.10).

Prey use was likewise unrelated to the natural prey availability of the corresponding guilds. Neither the main effect of relative prey availability (χ² = 0.01, df = 1, p = 0.90) nor its interaction with prey type (χ² = 1.22, df = 3, p = 0.75) significantly affected attack probability. Consistent with this, none of the prey-specific slopes differed significantly from zero (all p ≥ 0.35), and no pairwise differences among slopes were detected after Tukey adjustment (all p ≥ 0.73). Notably, green larvae remained the most frequently attacked prey type despite being among the least abundant in the environment, whereas ants were the most abundant natural prey guild but among the least attacked (**Fig. 4c**). Together, these results indicate selective prey use rather than tracking of local prey availability.

In contrast, prey use varied to some extent with the functional composition of the foraging bird assemblage (**Fig. 4d**). The interaction between prey type and the CWM of the proportion of invertebrates in the diet of foraging species was not significant overall (χ² = 5.19, df = 3, p = 0.16). Examined individually, however, attack probability on spider models tended to increase with this CWM, a trend that approached significance (β = 0.33 ± 0.18, p = 0.067), whereas no such relationship was found for ants (β = −0.19 ± 0.25, p = 0.45), brown larvae (β = −0.04 ± 0.18, p = 0.84), or green larvae (β = 0.15 ± 0.16, p = 0.34); slopes did not differ significantly among prey types after Tukey adjustement (all p ≥ 0.22). This model explained a substantial proportion of variance in prey use (marginal R² = 0.44), suggesting that the insectivory trait of species within foraging assemblages may shape prey selection, particularly for spiders.

Finally, bird species did not significantly differ in their use of prey types (χ² = 21.62, p = 0.88; Monte Carlo permutations; **Fig. S6**). Although statistical power was limited by the low number of attacks recorded for most bird species, green larvae dominated the diet across all species, suggesting that these preferences were broadly shared within the bird community.

## Discussion

Whether urbanization-induced changes in bird community composition ultimately translate into altered trophic interactions has remained largely unresolved. By integrating functional traits, prey availability, experimental prey models, and direct observations of foraging interactions, we show that this translation occurs through successive ecological filters rather than directly from community composition. Urbanization simultaneously restructures bird communities, modifies prey resources, and filters both the species generating trophic interactions and the prey they exploit, revealing mechanisms that remain undetected when ecosystem functioning is inferred from biodiversity patterns alone.

### Urbanization restructures both insectivorous bird communities and their arthropod prey resources

We showed that increasing urbanization simultaneously altered the composition and functional structure of insectivorous bird communities and the availability of three of their main arthropod prey guilds. Bird community diversity, functional dispersion, and dietary specialization declined with increasing impervious surface cover, confirming that urban environments progressively favor more generalist communities (Palacio, 2020; Souza et al., 2019). However, these negative effects were partly mitigated by urban vegetation. Tree diversity buffered species richness loss, likely by increasing habitat availability and resource heterogeneity (Ferenc et al., 2014), whereas canopy cover buffered declines in Shannon diversity and FDis, highlighting the importance of wooded habitats for maintaining functionally diverse insectivorous assemblages (Basile et al., 2025; Cristaldi et al., 2017).

Urbanization also modified arthropod prey availability. The decline of lepidopteran larvae is consistent with their sensitivity to host plant loss and habitat fragmentation in cities (Burghardt et al., 2009; Soga & Koike, 2012). The decline in tree-associated spiders may reflect their dependence on structurally complex wooded habitats (Melo et al., 2021), whereas many urban-tolerant spiders occur in more open habitats (Magura et al., 2010). In contrast, the stability of ants is consistent with their generalist and disturbance-tolerant ecology (Menke et al., 2011; Uno et al., 2010). Together, these guild-specific responses suggest that urbanization alters prey composition more strongly than overall prey abundance, potentially affecting prey quality and palatability rather than quantity alone (Raupp et al., 2010; Seress & Liker, 2015), with potential consequences for bird-mediated trophic interactions and the ecosystem services they support.

Interestingly, unlike bird communities, arthropod prey did not benefit from higher local vegetation cover, canopy cover, or tree diversity. This finding is consistent with a global meta-analysis by Fenoglio et al., (2020), which similarly found no significant effect of vegetation cover on the response of arthropod communities to urbanization across the studies compiled. Thus, habitat features promoting insectivorous bird diversity do not necessarily sustain their prey resources. Such an asymmetry in the effects of urbanization on those two trophic levels provides an indication that bird community composition alone may only lead to a partial representation of realized insectivory.

### Non-random ecological filtering decouples potential and realized insectivory

A central finding of this study is that the functional composition of insectivorous bird communities was insufficient to predict the trophic interactions they ultimately delivered (see also Schillé et al., 2025). Indeed, the subset of species observed foraging was consistently less insect-specialized than expected from the corresponding acoustically detected community. Null model simulations confirmed that this mismatch did not arise from random sampling of the plot species pool. Thus, bird community composition captures only the potential for insectivory, whereas realized ecosystem functions are determined by species that actually generate trophic interactions. This finding challenges a common assumption in trait-based ecology, that ecosystem functioning can be inferred directly from functional traits within communities (Palacio et al., 2018; Sol et al., 2020), implicitly assuming proportional contributions of all co-occurring species to the function. By using near-continuous camera systems to directly record foraging interactions, we moved beyond trait-based inference and characterizes realized trophic function more explicitly (Mrazova et al., *in prep*).

Several non-mutually exclusive mechanisms may underlie this potential-realized insectivory decoupling. First, generalist and synurbic species (e.g. House Sparrow) may disproportionately contribute to observed foraging interactions even when more insect-specialized species are present, reflecting reliance on opportunistic arthropod exploitation (Palacio, 2020), particularly under nutritional constraints linked to anthropogenic food (Bernat-Ponce et al., 2023). Their high abundance may further pre-empt arthropod resources from more specialized insectivores. Second, curiosity-driven and flexible behaviors may allow low-insectivorous species to exploit novel arthropod resources (Tryjanowski et al., 2016). Third, fine-scale habitat constraints, such as microhabitat structure, vegetation complexity, and substrate availability, can differentially affect foraging efficiency among co-occurring species (Shang et al., 2026; Whelan, 2001), producing unequal functional contributions not captured by community composition alone. Finally, artificial prey may not fully replicate natural prey cues (Zvereva & Kozlov, 2023); although this limitation likely applies consistently across plots, it could differentially affect attack rates if synurbic birds are more responsive to artificial stimuli than specialists.

Interestingly, the mismatch tended to decrease with increasing local vegetation cover, indicating closer alignment between realized and potential trophic interactions in more vegetated contexts, probably through a higher contribution of specialist species. Although not significant, plots showing the highest realized insectivory (23A, 19B, MNV; **Fig.1**) were characterized by structurally complex, dense, less managed vegetation with a well-developed understory, suggesting that metrics such as NDVI may not fully capture structural features relevant to foraging, particularly vertical complexity and microhabitat heterogeneity (Mitchell et al., 2016). Future studies should test whether finer structural descriptors better explain this translation. If landscape-scale features primarily determine bird community composition while fine-scale complexity shapes predation function (as observed for arthropods, Beninde et al., 2015), maintaining functionally diverse bird communities may be insufficient to sustain insectivory in cities, and small patches of structurally complex vegetation could play a complementary role (Nell et al., 2018).

### Prey use is associated with prey preferences and bird foraging assemblage composition rather than prey availability

Across the urban gradient, birds consistently preferred green caterpillar models over brown, while spiders and ants were attacked even less frequently. This ranking remained remarkably stable despite substantial variation in bird community composition and arthropod availability, indicating that prey preferences were largely conserved across urban contexts and likely reflect differences in prey profitability or quality (Razeng & Watson, 2015). None of the environmental predictors tested significantly altered prey preferences, suggesting that urbanization had a negligible influence on prey selection overall.

The observed prey preferences are unlikely to be explained solely by methodological artefacts. Contrary to previous studies reporting lower attack rates on low-luminance (e.g., brown) prey in temperate systems (Zvereva et al., 2019), green caterpillars were here consistently more attacked than brown ones, possibly reflecting differences in background matching against tree bark, while the smaller size of spider and ant models may have reduced their detectability. Nevertheless, because model color and size closely matched the observed natural prey, we interpret the results as ecologically realistic rather than methodological bias.

More surprisingly, these preferences were unrelated to the availability of the corresponding natural prey guilds. Lepidopteran larvae remained the most frequently attacked despite being among the least abundant arthropods across study plots, whereas ants were the most abundant prey but among the least attacked. Thus, realized trophic interactions may not be primarily driven by prey encounter rates and are probably instead consistent with selective prey use. Similar patterns have been reported in California, where nutritionally valuable dipterans and lepidopterans were disproportionately consumed by the Coastal Cactus Wren compared to other prey, despite being relatively less abundant in the environment (Nell et al., 2023).

Variation in prey use was rather associated with the functional composition of foraging communities. The CWM of the proportion of invertebrates in foraging birds’ diets emerged as an informative predictor, suggesting that realized prey use depends more on the functional groups engaged in trophic interactions than on environmental conditions or prey availability. This reinforces our finding that community composition alone is insufficient to predict realized trophic interactions: prey use also reflects the functional composition of the actively foraging assemblage. Insect-specialized communities tended to increase attacks on spiders, whereas no such relationship was detected for ants or brown caterpillars. Although marginal, this pattern is consistent with the high nutritional value of spiders for nestling development (Ramsay & Houston, 2003) and their importance early in the breeding season (Naef-Daenzer et al., 2000).

Finally, no significant variation in prey use was detected at the species level, potentially due to the limited number of predation events and statistical power. Nevertheless, selective exploitation of profitable prey might emerge as a community-level pattern rather than being driven solely by species traits in urban environments. Future studies using nest-box cameras or food-choice experiments would still be valuable for quantifying species-specific variation in prey choice under natural conditions. (Senar et al., 2021; Sinkovics et al., 2021).

## Conclusion

Because species present in a community are not necessarily those that generate ecological interactions, and prey encountered are not necessarily those consumed, distinguishing potential from realized interactions may be broadly relevant wherever functional inference is used to link animal communities to ecosystem processes.

In cities specifically, our findings suggest that conserving ecosystem functions requires more than maintaining biodiversity. While tree canopy cover and tree diversity offer simple levers for mitigating the negative effects of urbanization on insectivorous bird diversity, ensuring that this diversity translates into greater realized trophic functions may require additional habitat conditions, such as maintaining well-developed understory vegetation. As urban communities continue to reorganize under global change, understanding how novel species assemblages translate into trophic interactions will be critical for predicting the persistence of ecological functions such as natural pest control. Approaches that combine functional ecology with direct observations of species interactions therefore provide a promising framework for designing urban green spaces that support not only biodiversity, but also the ecological processes underpinning ecosystem functioning.

## Acknowledgement

This research was financially supported by the Natural Sciences and Engineering Research Council of Canada (NSERC) through an Alliance grant [ALLRP 571966-2022] to Alain Paquette. We also acknowledge Ville de Montréal (Service des grands parcs, du Mont-Royal et des sports, Service de l’eau, Arrondissements de Rosemont - La Petite Patrie and Ville-Marie), Ville de Dieppe, Ville de Repentigny, Soverdi, QuébecVert, Centre d’expertise et de recherche en infrastructures urbaines, Jakarto, Rousseau-Lefebvre, and Dominique Savio, for their financial and in-kind contributions as partner organizations in this Alliance grant.

We thank Matthieu Legay-Jeanjacquot, Matthias Briguet, Catherine Bérubé, Sarah Tardif, and Liliane Morin for their help in the field and we thank Charlotte Langlois, Arno Landry, Thania Veilleux-Gomez, and Pascale Bérubé for their contribution to sorting the photographs, as well as Emma Bacon, Marine Fernandez and Carly Ziter for the tree inventories. We thank Luc Barbaro, Bastien Castagneyrol, Anna Mrazova and Arndt Hampe for constructive discussions that helped improve the clarity and interpretation of our results.

During the preparation of this manuscript, two AI-assisted tools were used for language improvement exclusively: ChatGPT (OpenAI, GPT-5.2, 2026) and Grammarly (Grammarly Inc., 2024). ChatGPT was used to enhance clarity, consistency in language, and refine the academic writing style. Grammarly was employed to support grammar checking and spelling correction. All outputs from these tools were critically reviewed, edited, and approved by the authors to ensure accuracy, integrity, and originality.

## Author contribution

Laura Schillé conceived the ideas and designed the methodology. Frédéric Raspail, Philippe Chaumeil, Pierre Bordenave, and Laura Schillé designed the cameras and enabled the automation of image analysis. Laura Schillé, Vanessa Poirier, and Pierre-Alexis Herrault collected the data. Vanessa Poirier processed audio recordings for bird identification. Laura Schillé analyzed the data. Laura Schillé and Alain Paquette led the writing of the manuscript. Alain Paquette acquired the funding, provided the facilities, and supervised Laura Schillé. All authors contributed critically to the drafts and gave final approval for publication.

## Data availability statement

The statistical analyses were conducted using the R Quarto format to ensure full reproducibility. Both the code and the data will be made publicly available in accordance with the FAIR principles for scientific data management.

## Copyright statement

For Open Access, a CC-BY 4.0 public copyright license (https://creativecommons.org/licenses/by/4.0/) has been applied by the authors to the present document and will be applied to all subsequent versions up to the Author Accepted Manuscript arising from this submission.

