## Supplementary Materials S1 to S6 for "From bird communities to trophic interactions: successive ecological filters decouple potential and realized insectivory in urban environments"

*Appendix S1. Acoustic data processing, validation, and derivation of community metrics*

Recordings were conducted in the audible frequency range for 30 minutes per hour, from 30 minutes before sunrise to 30 minutes after sunset.

Audio files were processed using BirdNET (Kahl et al., 2021) through the *birdnetR* package, using the Meta v2.4 and TFLite v2.4 models in R (v. 4.4.2; R Core Team, 2021). Recordings were segmented into 3-second intervals and assigned to bird species that were geographically and temporally plausible, each associated with a confidence score. Consecutive detections of the same species separated by one second or less were grouped into a single vocalization event, assumed to originate from the same individual.

A subset of detections was validated by an experienced ornithologist (V.P.) using two randomly selected 10-minute segments per plot, taken from the morning chorus and balanced between weekdays and weekends to account for variation in anthropogenic noise. BirdNET identifications were classified as true positives (TP), false positives (FP), or false negatives (FN). Species-specific confidence thresholds were determined following Funosas et al. (2024). For species with at least 30 detections and at least 10 validated TP, thresholds were optimized using Youden's index (J = sensitivity + specificity − 1). For species not meeting these criteria, a conservative threshold of 0.9 was applied. Rare species absent from the validated subset were further checked manually using three randomly selected detections. The final dataset was obtained by retaining validated TP and FN while excluding FP.

From this dataset, we retained only functional insectivores (FI), defined as species known to consume arthropods and forage in trees during at least part of the year. This resulted in a total of 37 species across all plots.

To estimate FI species richness at the plot level, only species validated by the ornithologist were retained to minimize potential biases associated with automated detections. In contrast, all validated detections were used to quantify community structure. For each species, vocal dominance was calculated as the mean duration of its vocalizations divided by the total vocalization duration of all FI species within a plot. This metric was used to compute Shannon diversity and functional dispersion (FDis).

Functional traits (morphological, reproductive, and behavioral) were extracted from the Cornell Laboratory of Ornithology (2022) and EltonTraits 1.0 (Wilman et al., 2014) databases (**Table S1**). Functional dispersion was calculated using the *dbFD* function from the *FD* package (Laliberté et al., 2014).

Finally, we calculated a community-weighted mean (CWM) of the proportion of invertebrates in species' diets within the plot-level FI communities weighted by their vocal dominance. The proportion of invertebrates in each species' diet was derived from EltonTraits 1.0. Because arthropods constitute the dominant component of invertebrate prey consumed by FI birds, this metric represents *potential insectivory*, i.e., the expected contribution of the local bird community to arthropod consumption based on its composition.

**Eq. S1**

$$CWM_{\%\ invertebrates\ in\ diets,FI} = \sum_{i=1}^{N} v_i \times I_i$$

*$N$ is the number of species in the plot-level FI community,*
*$v_i$ is the relative vocal dominance of the species, calculated as its vocal dominance divided by the sum of vocal dominance across all FI species (that were acoustically detected at the plot level)*
*$I_i$ is the mean proportion of invertebrates in the diet of the species $i$ extracted from EltonTraits 1.0 database.*

*Table S1. Morphological, ecological, and life-history traits related to foraging and trophic ecology of the bird species included in the study (Cornell Laboratory of Ornithology, 2026; Wilman et al., 2014)*

| Genus species | Common Name | Habitat | Habitat Density | Nest | Eggs | Forag | Trophic Level | Trophic Niche | Primary Lifestyle | % of Invertebrates in diet | Beak Length Culmen | Mass | Conservation | Migration | Range Size |
|---|---|---|---|---|---|---|---|---|---|---|---|---|---|---|---|
| Agelaius phoeniceus | Red-winged Blackbird | Wetland | 3 | Shrub | [2-4] | Ground Forager | Herbivore | Granivore | Generalist | 50 | 22,7 | 50,8 | Low Concern | 3 | 13703226,95 |
| Bombycilla cedrorum | Cedar Waxwing | Woodland | 2 | Tree | [2-6] | Foliage Gleaner | Herbivore | Frugivore | Insessorial | 20 | 15,1 | 31,6 | Low Concern | 3 | 7437954,14 |
| Cardellina pusilla | Wilson's Warbler | Forest | 1 | Ground | [2-7] | Foliage Gleaner | Carnivore | Invertivore | Insessorial | 80 | 11,6 | 7 | Low Concern | 3 | 7961357,24 |
| Cardinalis cardinalis | Northern Cardinal | Shrubland | 2 | Shrub | [2-5] | Ground Forager | Herbivore | Omnivore | Insessorial | 20 | 17,6 | 42,6 | Low Concern | 1 | 5834572,06 |
| Catharus fuscescens | Veery | Forest | 1 | Ground | [1-5] | Ground Forager | Carnivore | Invertivore | Terrestrial | 50 | 17,6 | 31,9 | Low Concern | 3 | 3681872,46 |
| Catharus guttatus | Hermit Thrush | Forest | 1 | Ground | [3-6] | Ground Forager | Omnivore | Invertivore | Generalist | 80 | 18,8 | 30,1 | Low Concern | 3 | 8383811,92 |
| Certhia americana | Brown Creeper | Forest | 1 | Tree | [5-6] | Bark Forager | Carnivore | Invertivore | Insessorial | 70 | 16,2 | 8,1 | Low Concern | 1 | 6543644,5 |
| Coccyzus erythropthalmus | Black-billed Cuckoo | Forest | 1 | Tree | [2-5] | Foliage Gleaner | Carnivore | Invertivore | Insessorial | 60 | 27 | 50,9 | Common Bird in Steep Decline | 3 | 4944717,14 |
| Colaptes auratus | Northern Flicker | Shrubland | 2 | Cavity | [5-8] | Ground Forager | Carnivore | Invertivore | Generalist | 70 | 38,4 | 131,5 | Low Concern | 2 | 10804791,76 |
| Corvus brachyrhynchos | American Crow | Human Modified | 2 | Tree | [3-9] | Ground Forager | Omnivore | Omnivore | Terrestrial | 20 | 51,1 | 448,8 | Low Concern | 2 | 11573830,54 |
| Cyanocitta cristata | Blue Jay | Woodland | 2 | Tree | [2-7] | Ground Forager | Omnivore | Omnivore | Terrestrial | 20 | 29,9 | 88 | Low Concern | 1 | 6665383,16 |
| Dryobates pubescens | Downy Woodpecker | Woodland | 2 | Cavity | [3-8] | Bark Forager | Carnivore | Invertivore | Insessorial | 80 | 17,4 | 25,6 | Low Concern | 1 | 12800223,82 |
| Dryocopus pileatus | Pileated Woodpecker | Forest | 1 | Cavity | [3-5] | Bark Forager | Carnivore | Invertivore | Insessorial | 70 | 52,1 | 286,6 | Low Concern | 1 | 5925872,32 |
| Dumetella carolinensis | Gray Catbird | Shrubland | 1 | Shrub | [1-6] | Ground Forager | Omnivore | Invertivore | Insessorial | 60 | 19,2 | 35,3 | Low Concern | 3 | 6824562,12 |
| Empidonax flaviventris | Yellow-bellied Flycatcher | Forest | 1 | Ground | [2-5] | Flycatching | Carnivore | Invertivore | Insessorial | 90 | 12,9 | 11,8 | Low Concern | 3 | 5332665,34 |
| Geothlypis trichas | Common Yellowthroat | Wetland | 1 | Shrub | [1-6] | Foliage Gleaner | Carnivore | Invertivore | Insessorial | 100 | 12,7 | 9,5 | Low Concern | 3 | 12826786,25 |
| Hesperiphona vespertina | Evening Grosbeak | Forest | 1 | Tree | [2-5] | Ground Forager | Herbivore | Omnivore | Insessorial | 10 | 22,7 | 57,3 | Alert | 2 | 3942604,88 |
| Junco hyemalis | Dark-eyed Junco | Forest | 2 | Ground | [3-6] | Ground Forager | Herbivore | Granivore | Generalist | 30 | 11,6 | 19,5 | Low Concern | 3 | 10167654,77 |
| Leiothlypis ruficapilla | Nashville Warbler | Forest | 1 | Ground | [4-5] | Foliage Gleaner | Carnivore | Invertivore | Insessorial | 80 | 11,6 | 8,1 | Low Concern | 3 | 2772463,48 |

| | | | | | | | | | | | | | | | |
|---|---|---|---|---|---|---|---|---|---|---|---|---|---|---|---|
| Leuconotopicus villosus | Hairy Woodpecker | Forest | 1 | Cavity | [3-6] | Bark Forager | Carnivore | Invertivore | Insessorial | 70 | 31,6 | 62,7 | Low Concern | 1 | 13332874,44 |
| Melanerpes carolinus | Red-bellied Woodpecker | Woodland | 1 | Cavity | [2-6] | Bark Forager | Omnivore | Omnivore | Insessorial | 30 | 32,6 | 69,5 | Low Concern | 1 | 3074161,82 |
| Melospiza melodia | Song Sparrow | Shrubland | 2 | Shrub | [1-6] | Ground Forager | Omnivore | Omnivore | Generalist | 40 | 13,4 | 21,9 | Low Concern | 2 | 10313728,38 |
| Myiarchus crinitus | Great Crested Flycatcher | Woodland | 1 | Cavity | [4-8] | Flycatching | Omnivore | Invertivore | Insessorial | 60 | 22,7 | 32,1 | Low Concern | 3 | 5137539,15 |
| Passer domesticus | House Sparrow | Human Modified | 3 | Cavity | [1-8] | Ground Forager | Herbivore | Granivore | Terrestrial | 10 | 13,5 | 26,5 | Low Concern | 1 | 34407867,59 |
| Passerina cyanea | Indigo Bunting | Shrubland | 2 | Shrub | [3-4] | Foliage Gleaner | Herbivore | Granivore | Terrestrial | 70 | 11,7 | 14,7 | Low Concern | 3 | 5700834,65 |
| Pheucticus ludovicianus | Rose-breasted Grosbeak | Forest | 2 | Tree | [1-5] | Foliage Gleaner | Herbivore | Omnivore | Insessorial | 50 | 18,6 | 42 | Low Concern | 3 | 3980871,24 |
| Piranga olivacea | Scarlet Tanager | Forest | 1 | Tree | [3-5] | Foliage Gleaner | Carnivore | Invertivore | Insessorial | 80 | 18,8 | 28,2 | Low Concern | 3 | 2605301,09 |
| Poecile atricapillus | Black-capped Chickadee | Forest | 1 | Cavity | [1-13] | Foliage Gleaner | Omnivore | Invertivore | Insessorial | 60 | 10,3 | 10,8 | Low Concern | 1 | 9078514,18 |
| Quiscalus quiscula | Common Grackle | Wetland | 3 | Tree | [1-7] | Ground Forager | Carnivore | Omnivore | Terrestrial | 40 | 33 | 105,2 | Common Bird in Steep Decline | 2 | 8216640,27 |
| Regulus satrapa | Golden-crowned Kinglet | Forest | 1 | Tree | [3-11] | Foliage Gleaner | Carnivore | Invertivore | Insessorial | 100 | 9 | 6,2 | Low Concern | 3 | 6348089,15 |
| Sayornis phoebe | Eastern Phoebe | Forest | 2 | Building | [2-6] | Flycatching | Carnivore | Invertivore | Insessorial | 90 | 17,4 | 19,7 | Low Concern | 3 | 6246276,45 |
| Setophaga magnolia | Magnolia Warbler | Forest | 1 | Tree | [3-5] | Foliage Gleaner | Carnivore | Invertivore | Insessorial | 100 | 12,1 | 8,1 | Low Concern | 3 | 3661483,1 |
| Setophaga pensylvanica | Chestnut-sided Warbler | Forest | 2 | Shrub | [3-5] | Foliage Gleaner | Carnivore | Invertivore | Insessorial | 80 | 13,4 | 9,3 | Low Concern | 3 | 2422614,2 |
| Setophaga aestiva/petechia | Yellow Warbler | Shrubland | 1 | Shrub | [1-7] | Foliage Gleaner | Carnivore | Invertivore | Insessorial | 100 | 11,7 | 10,2 | Low Concern | 3 | 15552185,22 |
| Setophaga pinus | Pine Warbler | Forest | 2 | Tree | [3-5] | Bark Forager | Carnivore | Invertivore | Insessorial | 70 | 13,4 | 11,8 | Low Concern | 3 | 2073441,98 |
| Setophaga ruticilla | American Redstart | Forest | 1 | Tree | [1-5] | Foliage Gleaner | Carnivore | Invertivore | Insessorial | 80 | 12,3 | 8,2 | Low Concern | 3 | 6675504,95 |
| Setophaga striata | Blackpoll Warbler | Forest | 1 | Tree | [3-5] | Foliage Gleaner | Carnivore | Invertivore | Insessorial | 70 | 13,9 | 11,8 | Steep Decline | 3 | 6353206,91 |
| Sitta canadensis | Red-breasted Nuthatch | Forest | 1 | Cavity | [2-8] | Bark Forager | Omnivore | Omnivore | Insessorial | 50 | 14 | 9,8 | Low Concern | 3 | 7299100,07 |

| | | | | | | | | | | | | | | | |
|---|---|---|---|---|---|---|---|---|---|---|---|---|---|---|---|
| Sitta carolinensis | White-breasted Nuthatch | Forest | 1 | Cavity | [5-9] | Bark Forager | Omnivore | Omnivore | Insessorial | 50 | 19,1 | 21 | Low Concern | 1 | 8811748,35 |
| Spinus tristis | American Goldfinch | Forest | 1 | Shrub | [2-7] | Foliage Gleaner | Herbivore | Granivore | Insessorial | 10 | 11,5 | 12,8 | Low Concern | 1 | 7802444,22 |
| Spizella passerina | Chipping Sparrow | Forest | 2 | Shrub | [2-7] | Ground Forager | Omnivore | Omnivore | Generalist | 40 | 13,1 | 12,2 | Low Concern | 3 | 13081064,54 |
| Sturnus vulgaris | European Starling | Human Modified | 3 | Cavity | [3-6] | Ground Forager | Omnivore | Omnivore | Insessorial | 20 | 30,2 | 77,1 | Low Concern | 2 | 17082191,68 |
| Turdus migratorius | American Robin | Forest | 3 | Tree | [3-5] | Ground Forager | Omnivore | Invertivore | Generalist | 50 | 23,4 | 78,5 | Low Concern | 3 | 16455945,96 |
| Vireo gilvus | Warbling Vireo | Forest | 1 | Tree | [1-5] | Foliage Gleaner | Carnivore | Invertivore | Insessorial | 80 | 14,3 | 12,7 | Low Concern | 3 | 9680437,56 |
| Vireo olivaceus | Red-eyed Vireo | Forest | 1 | Tree | [1-5] | Foliage Gleaner | Omnivore | Invertivore | Insessorial | 60 | 16,4 | 16,1 | Low Concern | 3 | 8742854,76 |
| Vireo philadelphicus | Philadelphia Vireo | Woodland | 1 | Tree | [3-4] | Foliage Gleaner | Carnivore | Invertivore | Insessorial | 70 | 13,2 | 11,5 | Low Concern | 3 | 2750346,28 |

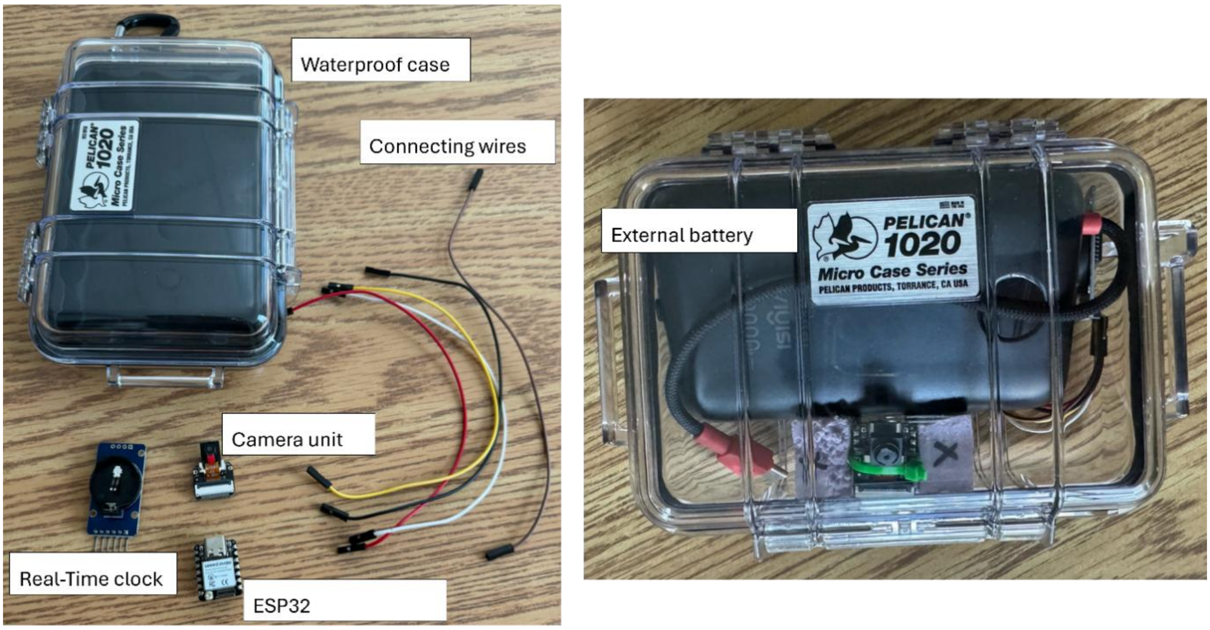


*Fig. S1. Architecture and components of the custom-built camera system deployed on trees to record bird foraging events. The system was based on ESP32 modules and camera sensors, powered by external batteries (20,000 mAh) and equipped with a real-time clock, all enclosed in a waterproof plastic case. Photographs taken by M. Briguet.*

## Eq. S2

$$CWM_{\%\ invertebrates\ in\ diets, foraging} = \sum_{i=1}^{n} o_i \times I_i$$

*$n$ is the number of species in the tree-level cafeteria foraging assemblage,*
*$o_i$ is the relative occurrence frequency of species $i$, calculated as its occurrence frequency divided by the sum of occurrence frequencies across all foraging species detected by the camera at the tree level*
*$I_i$ is the mean proportion of invertebrates in the diet of the species $i$ extracted from the EltonTraits 1.0 database.*

*Table S2. Summary of the environmental predictors of the study, their aggregation level, their definition, their resolution and their sources*

| Predictor | Aggregation level | Definition | Resolution | Source |
|---|---|---|---|---|
| Impervious surface | 200m around focal trees | Percentage of the combined surface area of low and high mineral surfaces | $1m^2$ | Normalized Difference Vegetation Index (NDVI) and city's digital elevation model (Communauté métropolitaine de Montréal, 2023) |
| Building cover | 200m around focal trees | Percentage of high mineral surface area (>3m) | $1m^2$ | Normalized Difference Vegetation Index (NDVI) and city's digital elevation model (Communauté métropolitaine de Montréal, 2023) |
| Canopy cover | 200m around focal trees | Percentage of high vegetation surface area (>3m) | $1m^2$ | Normalized Difference Vegetation Index (NDVI) and city's digital elevation model (Communauté métropolitaine de Montréal, 2023) |
| Low vegetation surface | 200m around focal trees | Percentage of low vegetation surface area (<3m) | $1m^2$ | Normalized Difference Vegetation Index (NDVI) and city's digital elevation model (Communauté métropolitaine de Montréal, 2023) |
| Local vegetation surface | 20m around focal trees | Percentage of the combined surface area of herbaceous, shrub, and tree layers | $1m^2$ | Normalized Difference Vegetation Index (NDVI) and city's digital elevation model (Communauté métropolitaine de Montréal, 2023) |
| Connectivity | 200m around focal trees | Hanski connectivity ($S_i$) between target trees i and each vegetation patch > 5m2<br>$$S_i = \sum A_j \times e^{-\propto \times d_{ij}}$$<br>With :<br>- $A_j$ : area of target patch j<br>- $\propto = \frac{1}{d}$ : parameter of connectivity decay with bird dispersal capacity ($d = 200m$)<br>- $d_{ij}$ : distance between tree i and patch j | $1m^2$ | Normalized Difference Vegetation Index (NDVI) and city's digital elevation model (Communauté métropolitaine de Montréal, 2023) |
| Population density | 200m around focal trees | Mean population density (persons.$km^{-2}$), computed as an area-weighted average of dissemination area population densities intersecting the 200m buffer. | Dissemination area | (Statistics Canada, 2022) |
| Nighttime radiance | 200m around focal trees | A proxy for light pollution. Mean nighttime light radiance ($nW \cdot cm^{-2} \cdot sr^{-1}$) from VIIRS DNB monthly cloud-free composites (EOG), June 2025. | ~450m | (Elvidge et al., 2013) |
| Surface temperature | 200m around focal trees | Daytime land surface temperature derived from airborne thermal infrared imagery, acquired during peak heating hours (12:00–15:00), August–September 2016. | 2m | (Ville de Montréal, 2016) |
| Biophony to anthropophony ratio | Plot level | A proxy for noise pollution (inversely proportional to anthropogenic noise pollution). NDSI index, calculated from passive acoustic field recordings. | ~200m plot level | Field data acquisition |
| Neighbor tree richness | Plot level | All public and private trees within a 200 m radius around the center of each plot were inventoried and identified to species level. | 200m plot level | (Paquette et al., 2026) |
| Neighbor tree Shannon diversity index | Plot level | Tree species diversity was quantified using a Shannon index weighted by basal area. Basal area (BA, $cm^2$) was calculated for each tree from DBH ($BA = \pi \times (\frac{DBH}{2})^2$. Trees with missing DBH values (1.6%) were excluded. | 200m plot level | (Paquette et al., 2026) |

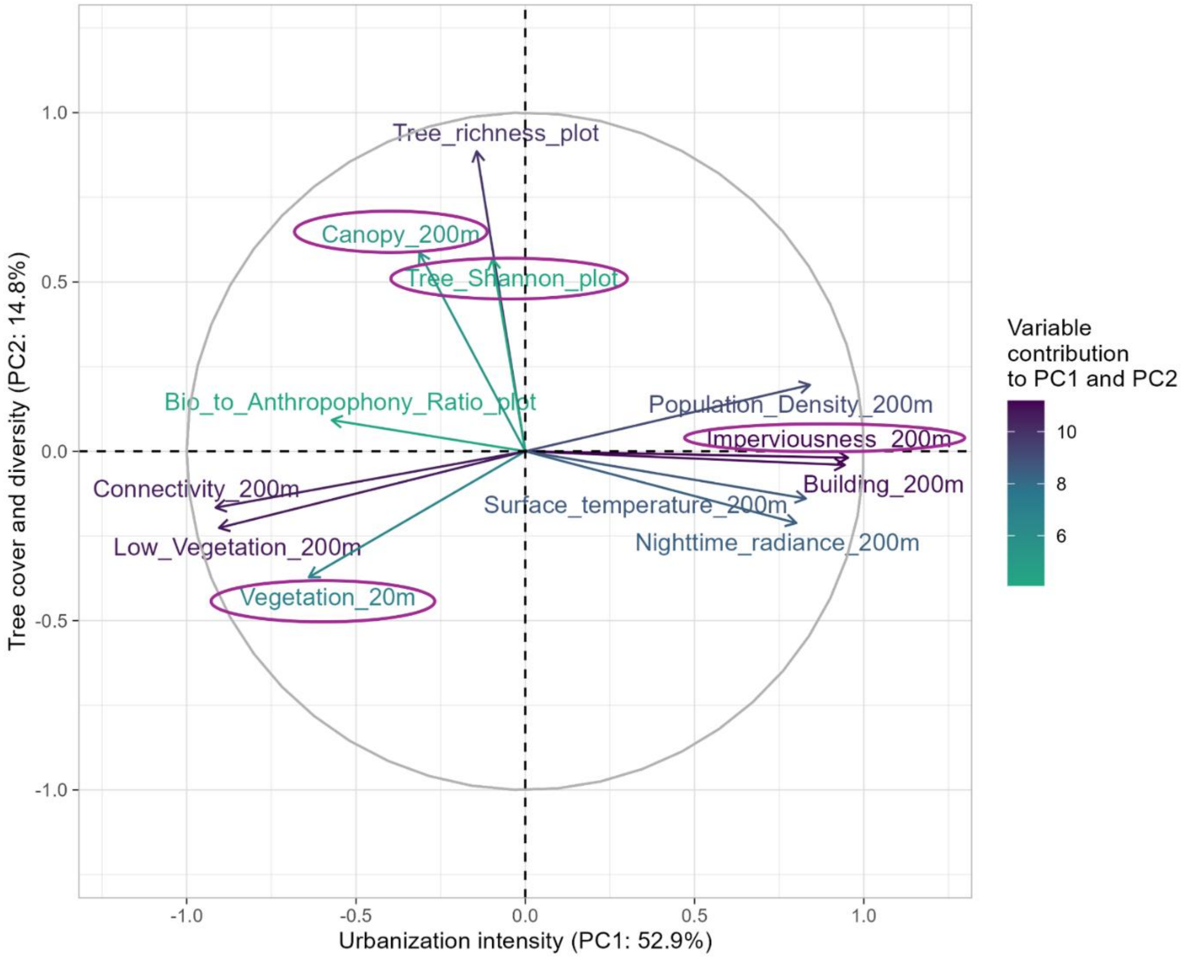


*Fig. S2. Principal component analysis summarizing the 12 tested environmental predictors. PC1 represents urban-related predictors and PC2 represents surrounding tree diversity. Variables retained for subsequent analyses are highlighted by a purple outline.*

*Table S3. Overview of statistical models used to test hypotheses H1–H3. The table summarizes response variables, model structures, fixed and random effects, key parameter estimates, and overall model fit. For models involving an interaction with prey type, Key results reports the likelihood-ratio test (LRT, via null-model comparison) for the main and interaction effects, together with prey-specific slopes and Tukey-adjusted pairwise comparisons obtained from emtrends. Significance levels: *** p < 0.001, ** p < 0.01, * p < 0.05, . p < 0.1. Arrows (↑/↓) indicate effect direction where a term reaches at least p < 0.1.*

| Hyp | Response variable | Model and distribution | Fixed effects | Random effects | Key results and associated statistics | Statistics of the whole model |
|---|---|---|---|---|---|---|
| H1 | FI Specific richness | GLM Poisson (log) | Imperv_200m × (Canopy_200m + Tree_Diversity_plot) | N/A | Imperv_200m ↓*** (β± SE= -0.33 ± 0.10 ; z = −3.42 , $p < 0.001$)<br><br>Imperv_200m × Tree_Div _plot ↑. (β=0.21 ± 0.12; z = 1.75, p = 0.079) | LRT $\chi^2(5) = 17.10$, p = 0.004;<br>Pseudo-$R^2$ : 0.69 |
| H1 | FI Shannon diversity | LM | Imperv_200m × (Canopy_200m + Tree_Diversity_plot) | N/A | Imperv_200m ↓** (β=-0.35 ± 0.11; t = -3.15, $p < 0.01$)<br><br>Imperv_200m × Canopy_200m ↑* (β=0.21 ± 0.09; t = 2.24, $p < 0.05$) | $F(5,19) = 5.04$ , p = 0.004;<br>$R^2$adj = 0.46 |
| H1 | FI FDis | LM (sqrt transformation) | Imperv_200m × (Canopy_200m + Tree_Diversity_plot) | N/A | Imperv_200m ↓* (β=-0.037 ± 0.016; t = -2.32, $p < 0.05$)<br><br>Canopy_200m ↑. (β=0.027 ± 0.015; t = 1.82, p = 0.084)<br><br>Imperv_200m × Canopy_200m ↑* (β=0.034 ± 0.013; t = 2.59, $p < 0.05$) | $F(5,19)=5.26$, p = 0.003 ;<br>$R^2$adj = 0.47 |
| H1 | CWM % of invertebrates in diet of the FI community | LM (sqrt transformation) | Imperv_200m × (Canopy_200m + Tree_Diversity_plot) | N/A | Imperv_200m ↓*** (β=-0.84 ± 0.16 ; t = −5.2 , $p < 0.001$) | $F(5,19)=7.35$, p = 0.0005 ;<br>$R^2$adj = 0.57 |
| H1 | Natural prey guild abundance | Zero-inflated negative binomial (ZINB) | Real_Prey_Type × Imperv_200m | Plot | Ants > Spiders > Larvae (overall) *** (emmeans pairwise, all $p < 0.0001$)<br><br>Real_Prey_Type × Imperv_200m (overall) * (LRT $\chi^2(2) = 6.45$, p = 0.040)<br><br>Larvae ↓ Imperv_200m (slope) * (β = −0.56 ± 0.28, z = −2.02, p = 0.044)<br><br>Spiders ↓ Imperv_200m (slope) * (β = −0.51 ± 0.25, z = −2.02, p = 0.043)<br><br>Ant (slope ns) (β = 0.12 ± 0.20, p = 0.559)<br><br>No pairwise differences among slopes after Tukey (all $p \geq 0.087$).<br><br><u>NB</u>. Interactions with Vegetation_20m; Canopy200m; and Tree_Diversity_plot were not significant | LRT $\chi^2(5) = 103.5$, $p < 0.001$; $R^2$m = 0.49; $R^2$c = 0.54 |

| | | | | | | |
|---|---|---|---|---|---|---|
| H2 | Functional mismatch (Realized – Potential insectivory) | Wilcoxon signed-rank test (H0: median = 0) | N/A | N/A | Median mismatch = -6.10<br>(foragers less insectivorous than expected) | Wilcoxon signed-rank test; $V = 377$; $p < 0.01$ |
| H2 | Non-random assembly of foraging assemblages (null model test on SES) | One-sample t-test (H0: SES = 0) | N/A | N/A | mean SES = −0.51 ± 0.14 (foraging assemblages less insect-specialized than expected under random assembly) | One-sample t-test; $t = -3.70$; $p < 0.001$ |
| H2 | Functional mismatch (Realized – Potential insectivory) | LMM | Vegetation_20m | Plot | Vegetation_20m ↑ (ns)<br>($\beta = 0.097 \pm 0.077$; $t = 1.26$, $p = 0.214$)<br><br>NB. Imperv_200m, Canopy_200m and Tree_Diversity_plot were also tested and not significant | LRT $\chi^2(1) = 1.65$, $p = 0.199$; $R^2m = 0.037$; $R^2c = 0.415$ |
| H3 | Probability of attack | GLMM binomial (logit) | Prey_Type | Plot/Tree + Date | P(Attack): Green Larvae > Brown Larvae > Ants ≈ Spiders<br>(Tukey: all pairwise $p < 0.01$ except Ant vs. spider, $p = 0.79$) | LRT $\chi^2(3) = 123.25$, $p < 0.001$ ; $R^2m = 0.09$ ; $R^2c = 0.26$ |
| H3 | Probability of attack | GLMM binomial (logit) | Prey_Type × Imperv_200m | Plot/Tree + Date | Prey_Type × Imperv_200m (overall ns)<br>(LRT $\chi^2(3) = 3.85$, $p = 0.28$)<br><br>No prey-specific slope differs from zero (all $p \geq 0.27$)<br><br>No pairwise differences after Tukey (all $p \geq 0.26$) | LRT $\chi^2(7) = 127.37$, $p < 0.001$ ;<br>$R^2m = 0.09$ ; $R^2c = 0.26$ |
| H3 | Probability of attack | GLMM binomial (logit) | Prey_Type × Tree_Diversity_plot | Plot/Tree + Date | Prey_Type × Tree_Diversity_plot (overall ns)<br>(LRT $\chi^2(3) = 4.33$, $p = 0.23$)<br><br>No prey-specific slope differs from zero (all $p \geq 0.13$)<br><br>No pairwise differences after Tukey (all $p \geq 0.18$) | LRT $\chi^2(7) = 127.85$, $p < 0.001$ ;<br>$R^2m = 0.10$ ; $R^2m = 0.27$ |
| H3 | Probability of attack | GLMM binomial (logit) | Prey_Type × Canopy_200m | Plot/Tree + Date | Prey_Type × Canopy_200m (overall ns)<br>(LRT $\chi^2(3) = 1.66$, $p = 0.65$)<br><br>No prey-specific slope differs from zero (all $p \geq 0.10$)<br><br>No pairwise differences after Tukey (all $p \geq 0.71$) | LRT $\chi^2(7) = 127.71$, $p < 0.001$;<br>$R^2m = 0.10$; $R^2c = 0.27$ |
| H3 | Probability of attack | GLMM binomial (logit) | Prey_Type × Vegetation_20m | Plot/Tree + Date | Prey_Type × Vegetation_20m (overall ns)<br>(LRT $\chi^2(3) = 2.51$, $p = 0.47$)<br><br>No prey-specific slope differs from zero (all $p \geq 0.30$)<br><br>No pairwise differences after Tukey (all $p \geq 0.43$) | LRT $\chi^2(7) = 125.76$, $p < 0.001$ ;<br>$R^2m = 0.09$ ; $R^2c = 0.27$ |
| H3 | Probability of attack | GLMM binomial (logit) | Prey_Type × Relative_Prey_availability | Plot/Tree + Date | Relative_Prey_availability (overall ns)<br>(LRT $\chi^2(1) = 0.01$, $p = 0.90$)<br><br>Prey_Type × Relative_Prey_availability (overall ns)<br>(LRT $\chi^2(3) = 1.22$, $p = 0.75$) | LRT $\chi^2(7) = 124.49$, $p < 0.001$; Marginal $R^2 = 0.09$ ; Conditional $R^2 = 0.26$ |

| | | | | | | |
|---|---|---|---|---|---|---|
| | | | | | No prey-specific slope differs from zero (all $p \geq 0.35$)<br><br>No pairwise differences after Tukey (all $p \geq 0.73$).<br><br><u>NB</u>.Green Larvae remained most attacked despite low availability; Ants most available but among least attacked | |
| H3 | Probability of attack | GLMM binomial (logit) | Prey_Type × CWM_invertDiet_of_forag | Plot | Prey_Type × CWM_invertDiet_of_forag (overall ns) (LRT $\chi^2(3) = 5.19$, $p = 0.16$)<br><br>P(Attack) on Spiders ↑ CWM_invertDiet_of_forag **.** ($\beta$=0.33 ± 0.18, $z = 1.83$, $p = 0.067$)<br><br>No other slope differs from zero (Ant $p = 0.45$; Brown Larvae $p = 0.84$; Green Larvae $p = 0.34$)<br><br>No pairwise differences after Tukey (all $p \geq 0.22$) | LRT $\chi^2(7) = 63.7$, $p <$ 0.001 ;<br>$R^2m = 0.44$ ; $R^2m = 0.92$ |
| H3 | Prey-type selection among bird species | Chi-square test | N/A | N/A | Did not differ significantly among bird species<br>($\chi^2 = 21.6$, $p = 0.88$, Monte-Carlo simulation)<br>Green larvae dominate the diet of all species (**Fig. S6**). | |

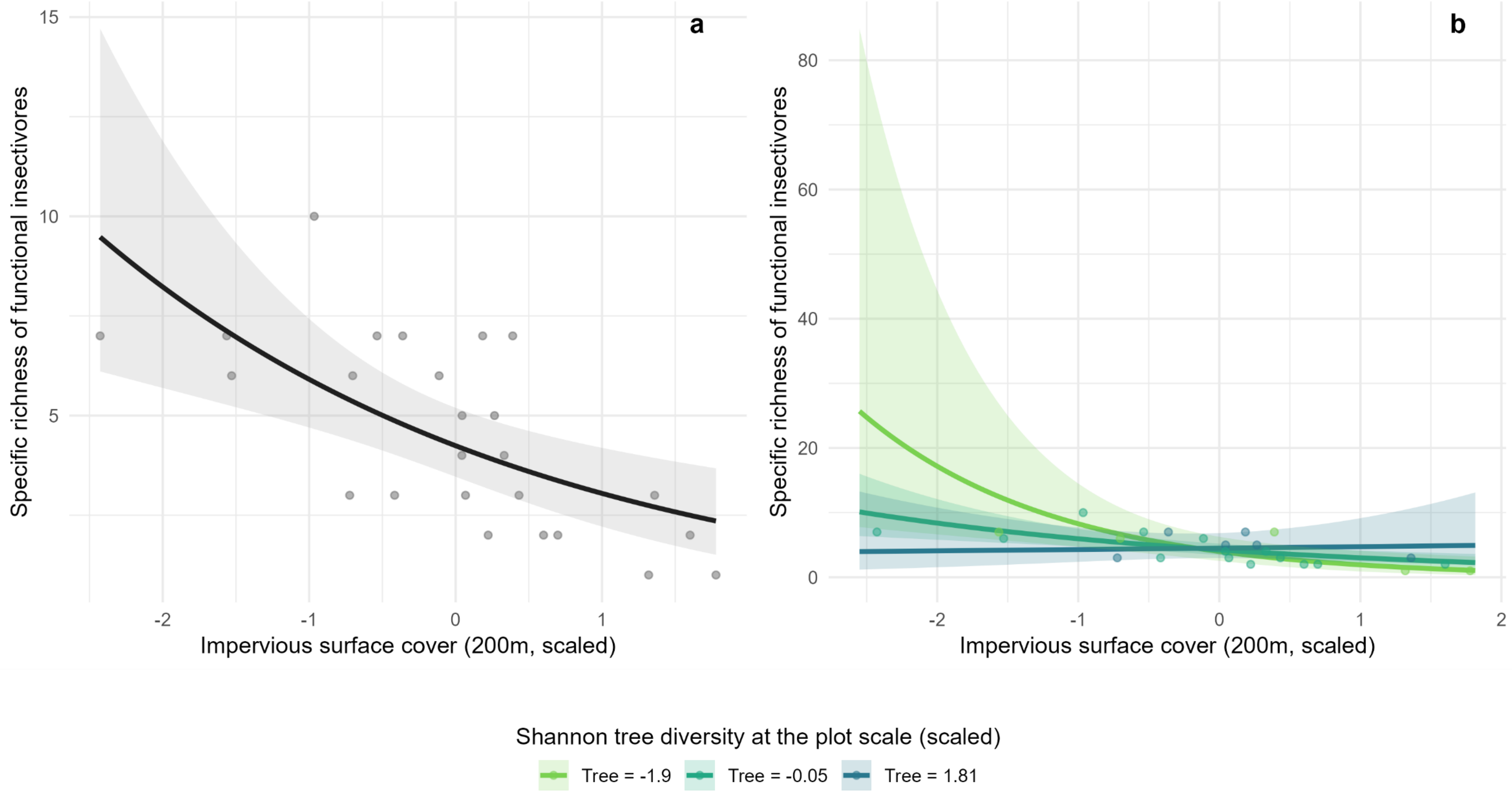


*Fig. S3. Predicted effects of impervious surface cover alone (a) and in interaction with plot-level tree diversity (b) on FI specific richness. Colored lines represent model predictions at low, intermediate, and high tree diversity, with shaded areas indicating 95% confidence intervals. Dots show raw data. Observed values for impervious surface ranged from 8.5 to 88.7% within a 200m buffer. Observed values for Shannon tree diversity ranged from 1.9 to 3.1 at the plot level.*

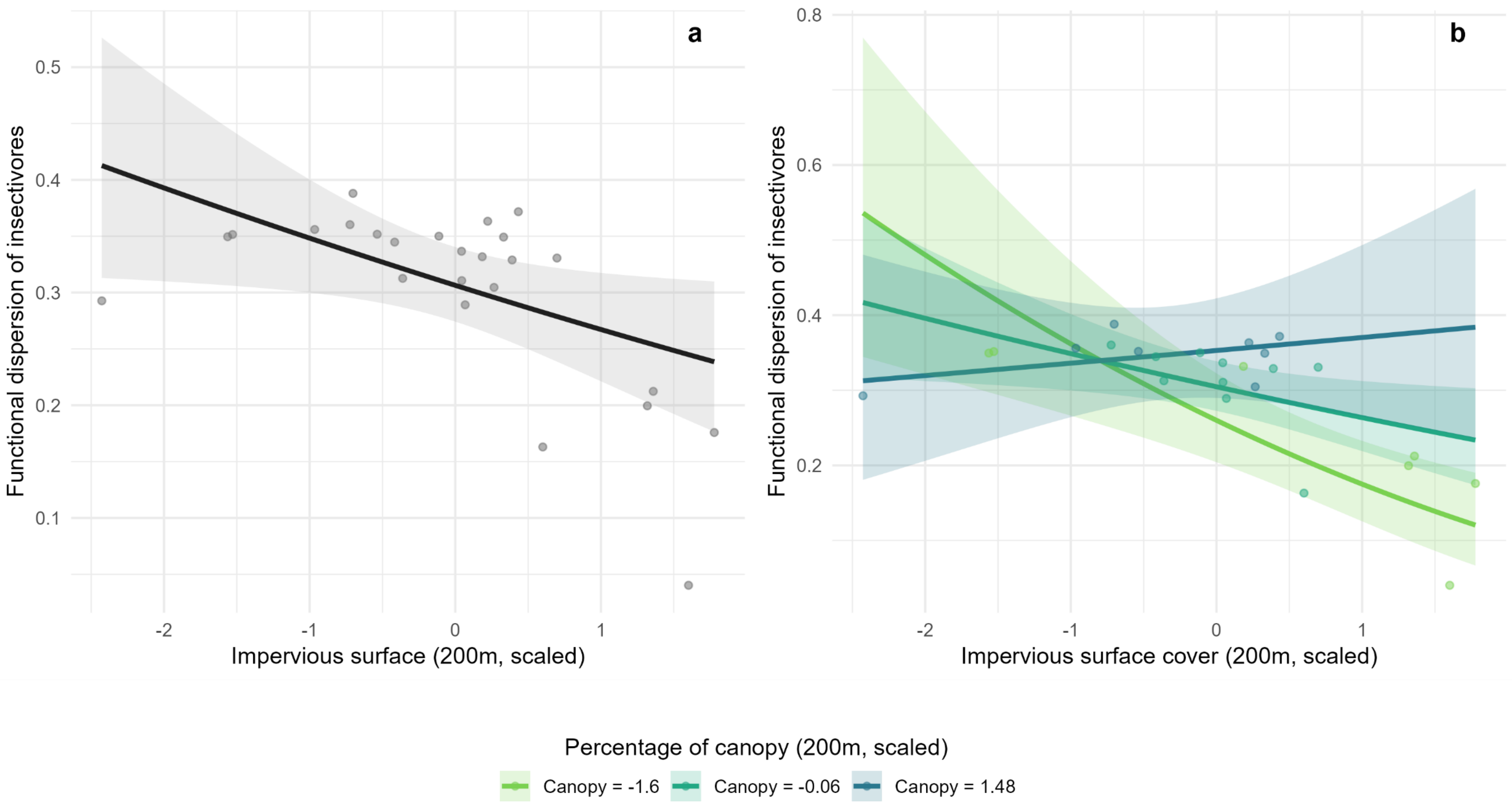


*Fig. S4. Predicted effects of impervious surface cover alone (a) and in interaction with canopy cover within 200m (b) on FI FDis. Colored lines represent model predictions at low, intermediate, and high canopy cover, with shaded areas indicating 95% confidence intervals. Dots show raw data. Observed values for impervious surface ranged from 8.5 to 88.7% within a 200m buffer. Observed values for canopy cover ranged from 8.8 to 32.0 within a 200m buffer.*

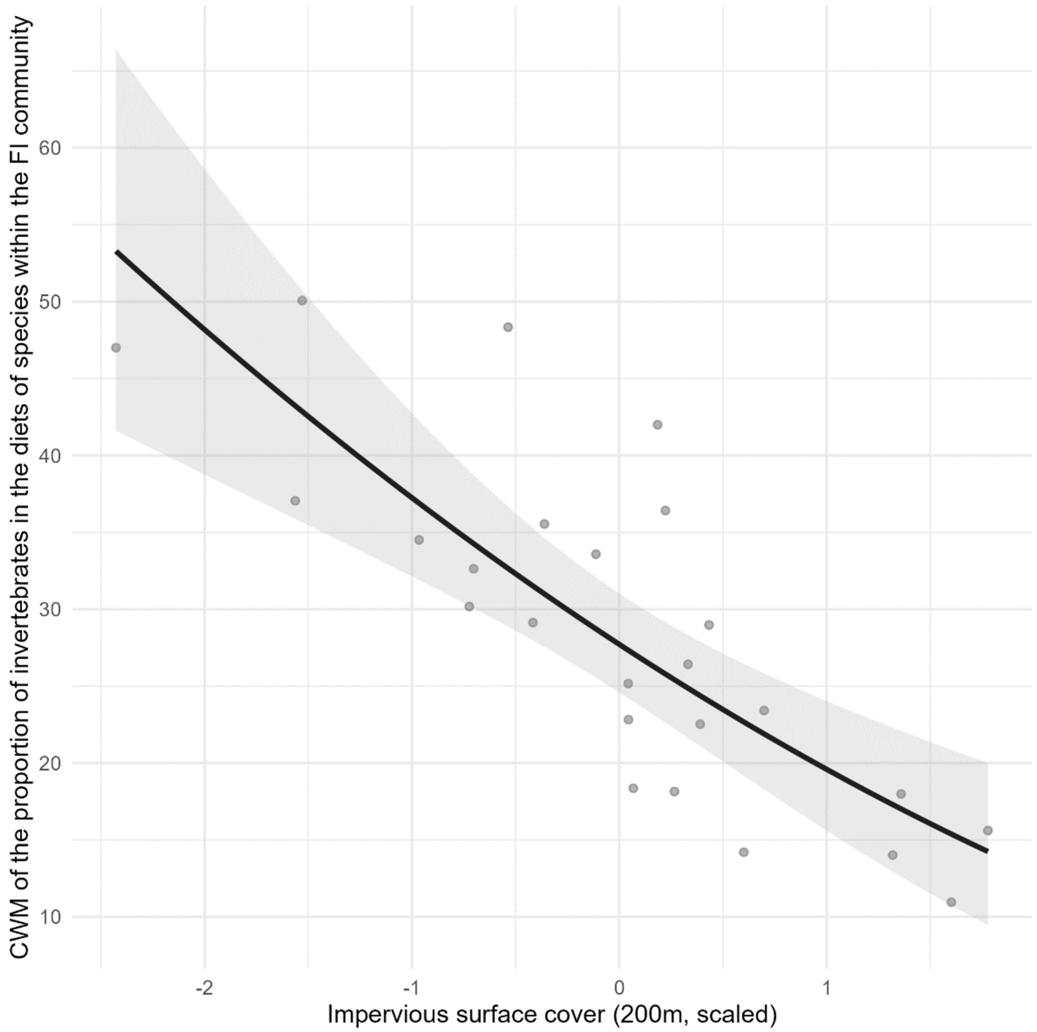


*Fig. S5. Predicted effects of impervious surface cover on CWM of the proportion of invertebrates in species' diet within the FI community. The shaded area indicates 95% confidence intervals. Dots show raw data. Observed values for impervious surface ranged from 8.5 to 88.7% within a 200m buffer.*

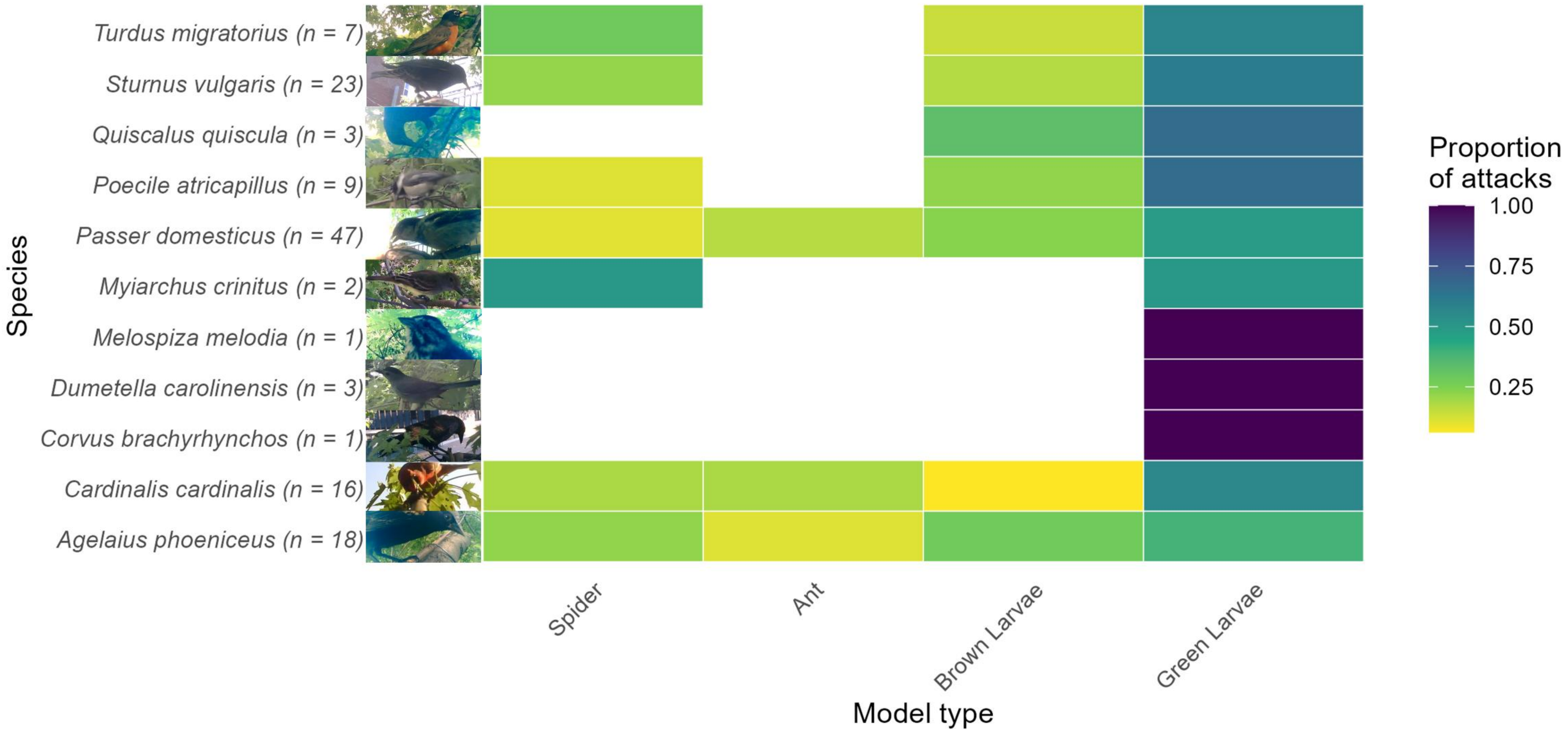


*Fig. S6. Proportion of attacks on four model prey types (spiders, ants, brown larvae, green larvae) by different bird species.*
